# Evolutionary genomics of socially polymorphic populations of *Pogonomyrmex californicus*

**DOI:** 10.1101/2021.03.21.436260

**Authors:** Mohammed Errbii, Ulrich R. Ernst, Aparna Lajmi, Eyal Privman, Jürgen Gadau, Lukas Schrader

**Author notes:** ^ǂ^equal contribution (Jürgen Gadau, Lukas Schrader). Apicultural State Institute, University of Hohenheim, Erna-Hruschka-Weg 6, DE-70599 Stuttgart, Germany. Center for Biodiversity and Integrative Taxonomy (KomBioTa), University of Hohenheim, DE-70599 Stuttgart, Germany. Corresponding authors: Lukas Schrader, Jürgen Gadau, **Email:**.

## Abstract

Social insects vary considerably in their social organization both between and within species. In the California harvester ant, *Pogonomyrmex californicus* (Buckley 1867), colonies are commonly founded and headed by a single queen (haplometrosis, primary monogyny). However, in some populations in California (USA), unrelated queens cooperate not only during founding (pleometrosis) but throughout the life of the colony (primary polygyny). The genetic architecture and evolutionary dynamics of this complex social niche polymorphism (haplometrosis vs pleometrosis) have remained unknown. Here, we provide a first analysis of its genomic basis and evolutionary history using population genomics comparing individuals from a haplometrotic population to those from a pleometrotic population. We discovered a recently evolved (< 200 k years), 8 Mb non-recombining region segregating with the observed social niche polymorphism. This region shares several characteristics with supergenes underlying social polymorphisms in other socially polymorphic ant species. However, we also find remarkable differences from previously described social supergenes. Particularly, four additional genomic regions not in linkage with the supergene show signatures of a selective sweep in the pleometrotic population. Within these regions, we find for example genes crucial for epigenetic regulation via histone modification (*chameau*) and DNA methylation (*Dnmt1*). These results suggest that social morph in this species is a polygenic trait involving a potential young supergene. Further studies targeting haplo- and pleometrotic individuals from a single population are however required to conclusively resolve whether these genetic differences underlie the alternative social phenotypes or have emerged through genetic drift.

## Introduction

Colonies of social Hymenoptera are led by a single (monogyny) or multiple reproductive females (polygyny) [1–3]. Such variations of social structure are suggested to have an adaptive value in certain conditions [2,4], and are conceptualized as alternative “social niches”, i.e. “the set of social environments in which the focal individual has non-zero inclusive fitness” [5]. In ants, species are often considered to be either monogyne or polygyne, but more and more cases of intraspecific variability in social organization have been described within several ant genera. Examples include multiple *Formica* and *Myrmica* species [6], various *Solenopsis* species [7–9], *Messor pergandei* [10], *Pogonomyrmex californicus* [11], *Cataglyphis niger* [12] *Leptothorax acevorum* [13] and *L. longispinosus* [14].

Species showing intraspecific variation in social organization are ideal to study the genomic architecture and evolutionary dynamics underlying complex trait polymorphisms. Recent population genomic studies in *Solenopsis* fire ants [9,15] and *Formica* wood ants [16–18] have provided significant insights into the origin, evolution, and geographic distribution of social variants in these species. Importantly, these studies led to the discovery of convergently evolved supergenes, i.e. genomic regions containing clusters of linked loci [19], underlying social niche polymorphisms in these species. Suppression of recombination in these supergenes drives the evolution of two or more diverged haplotype groups, similar to the X and Y haplotypes of sex chromosomes. Such genomic architecture facilitates the evolution and maintenance of complex trait polymorphisms (e.g. Brelsford et al., 2020; Hohenlohe et al., 2010; Joron et al., 2006; Matschiner et al., 2021; Nadeau et al., 2012). However social polymorphism can involve mechanisms other than supergenes as shown recently for the big-headed ant *Pheidole pallidula*, where genetic and environmental factors are suspected to contribute to the expression of alternative social morphs [24].

The California harvester ant *Pogonomyrmex californicus* [25] occurs throughout the Southwestern USA and Northwestern Mexico [26] and varies in terms of monogyny and polygyny, with primary monogyny as the predominant social organization [27]. In contrast to *Solenopsis* and *Formica* species, this variation is based on differences in colony founding strategy, i.e. haplometrosis (queens found a colony alone, “primary monogyny”) vs pleometrosis (multiple queens found a colony together, “primary polygyny”) [11,27–29], and not re-adoption of queens or colony budding [30,31]. Populations of *P. californicus* exhibit geographical variation in colony-founding strategies and are almost fixed for either the haplo- or the pleometrotic strategy [11,32]. Common garden experiments with foundresses belonging to these populations revealed that haplo- and pleometrotic behaviors are expressed independent of environmental conditions, consistent with a genetic polymorphism underlying these alternative social strategies [28,33], and representing adaptations to different ecological and geographic niches in this species.

In this study, we analyzed the genomic architecture, population dynamics, and evolutionary history of the social niche polymorphism in *P. californicus*, by comparing individuals from two populations in Southern California that are virtually fixed for either haplometrosis (H-population) or pleometrosis (P-population) [28,32]. We identify a non-recombining region of ca. 8 Mb that segregates with the studied populations, resembling previously described supergenes driving social polymorphisms in ants (Wang et al. 2013, Purcell et al. 2014). Additionally, we also find signatures of selective sweeps in the pleometrotic population outside the non-recombining region, suggesting that the social niche polymorphism in *P. californicus* is a polygenic trait involving not only a potential young supergene, but also additional unlinked modifier loci.

## Results

### Pogonomyrmex californicus genome assembly

To improve the available fragmented genome assembly, we combined minION long-read sequencing and 10X sequencing [34] of pleometrotic individuals to produce a highly contiguous reference genome assembly for *P. californicus*. The assembly spans 252.3 Mb across 199 scaffolds (N50 = 10.4 Mb, largest scaffold 20.75 Mb). We recovered 98.5% complete BUSCOs (S:97.8%,D:0.7%,F:1.1%,M:0.4%,n:4415), suggesting a nearly complete genome assembly. Genome annotation identified 22.79% repetitive sequences and 15,899 protein coding genes, resembling repeat and gene contents reported for other published ant genomes [35] (Dataset S1).

### The haplometrotic and pleometrotic populations are genetically distinct

For our population genomic analyses, we performed whole genome sequencing of 35 founding queens (19 from the P-population and 16 from the H-population). Principal Component Analyses (PCA) using 314,756 SNPs clearly separated queens of the two populations (Figures 1A and S1). Further, the PCA suggested higher genetic diversity in the H-population compared to the P-population. One founding queen (P99P) collected from the P-population was positioned between both populations in the PCA (Figure 1A), likely representing an F1 hybrid between a haplo- and a pleometrotic individual.

**Figure 1.**
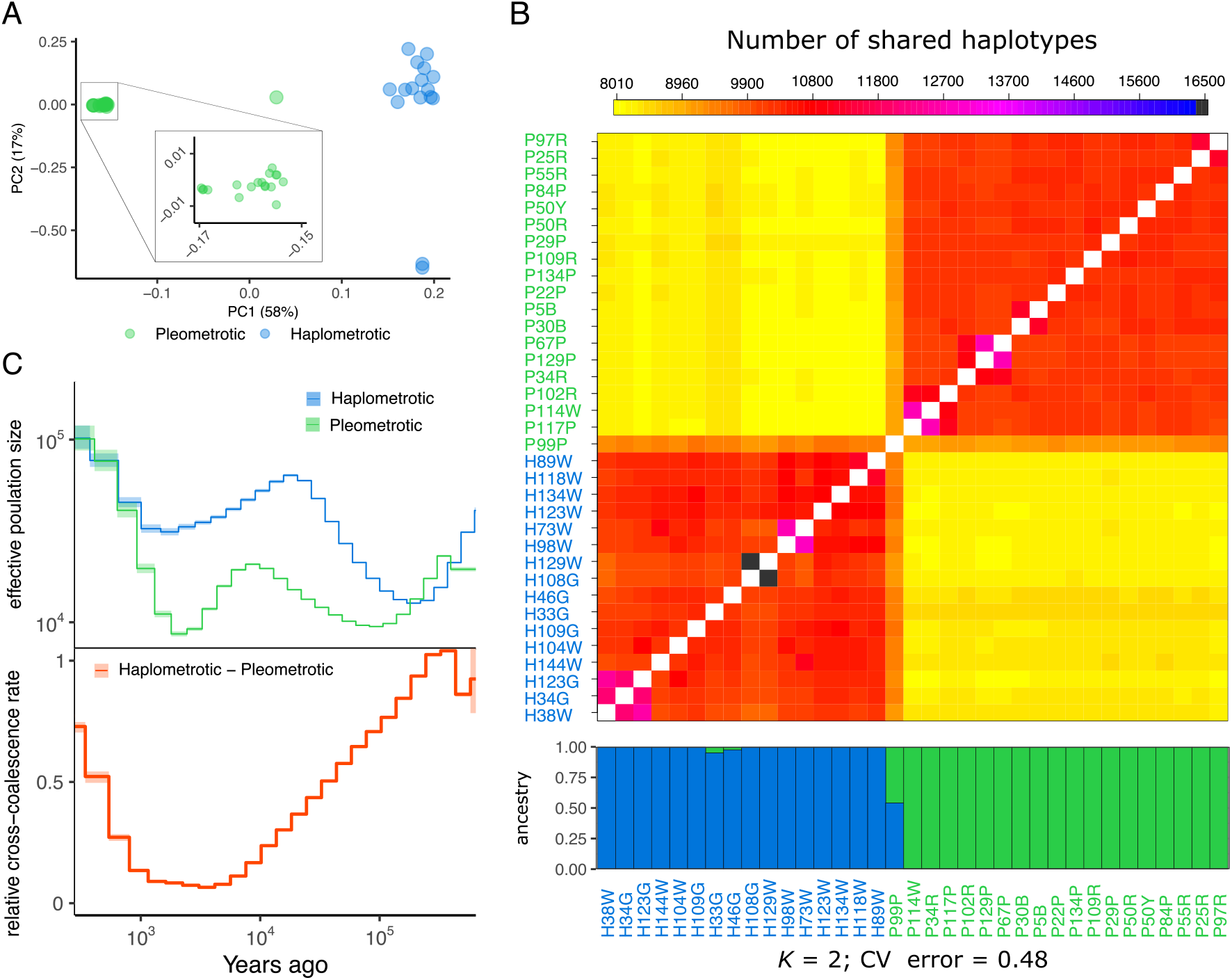
Population structure and demographic history in *P. californicus* populations. (A) Principal Component Analysis based on genome-wide SNP data of queens of the pleometrotic and haplometrotic populations (a scatterplot matrix showing four principal components can be found in Figure S1). (B) fineSTRUCTURE coancestry heatmap showing the number of shared haplotypes between donor (columns) and recipient (rows) queen pairs. ADMIXTURE plot below the heatmap shows patterns of ancestry among populations of *P. californicus* at k=2 (additional k values can be found in Figure S2). Note that one queen (P99P) is an apparent hybrid between the two populations. (C) Demographic history in *P. californicus* computed using MSMC2 with eight phased haplotypes (i.e. four queens), representing the pleometrotic and haplometrotic populations. Effective population size (*N*_e_) is displayed in the top panel and the relative cross-coalescence rate (rCCR) in the lower panel. Lines and shaded areas are means and 95% confidence intervals, respectively.

A detailed analysis of coancestry showed that queens share more haplotypes in within-population pairs than in between-population pairs (Figure 1B). Consistent with ongoing admixture between the two populations, we found significant between-population haplotype-sharing in some individuals (e.g. P99P, H33G and H46G) corroborated by ADMIXTURE analyses (Figures 1B and S2).

### Gene flow between the H- and P-populations is a recent phenomenon

We used MSMC2 [36] to reconstruct the demographic history of the P- and H-populations (Figure 1C). Both populations showed a similar trajectory with a zigzag-like pattern in effective population size (*N*_e_) between approximately 500,000 to 400 years ago, indicative of two separate bottlenecks: an ancient one about 400,000 until about 150,000 years ago and a recent one between 15,000 to roughly 2,000 years ago. Following the first bottleneck, *N_e_* consistently remained smaller in the pleometrotic population until between 2,000 and 400 years ago, when *N*_e_ started to increase again in both populations, reaching similarly high estimates in recent times.

We also inferred how these populations separated over time by estimating relative cross-coalescence (rCCR) [37] and found that the two populations started diverging during the first bottleneck at ∼300,000 years ago (Figure1C). Around 5,000 years ago, both populations were almost completely isolated (rCCR ≈ 0). However, until 400 years ago, rCCR increased rapidly, suggesting a growing genetic exchange, coinciding with population expansions in both populations. This predicted recent gene flow is corroborated by evidence of recent genetic admixture (Figures 1A and 1B).

### Genetic diversity is significantly lower in the pleometrotic population

The P-population is genetically less variable than the H-population based on genome-wide nucleotide diversity (𝜋) and heterozygosity (Figures 2A and 2B). Both estimates were significantly lower in the P-(median 𝜋 = 4.71e-4, median het = 0.167) compared to the H-population (median 𝜋

**Figure 2.**
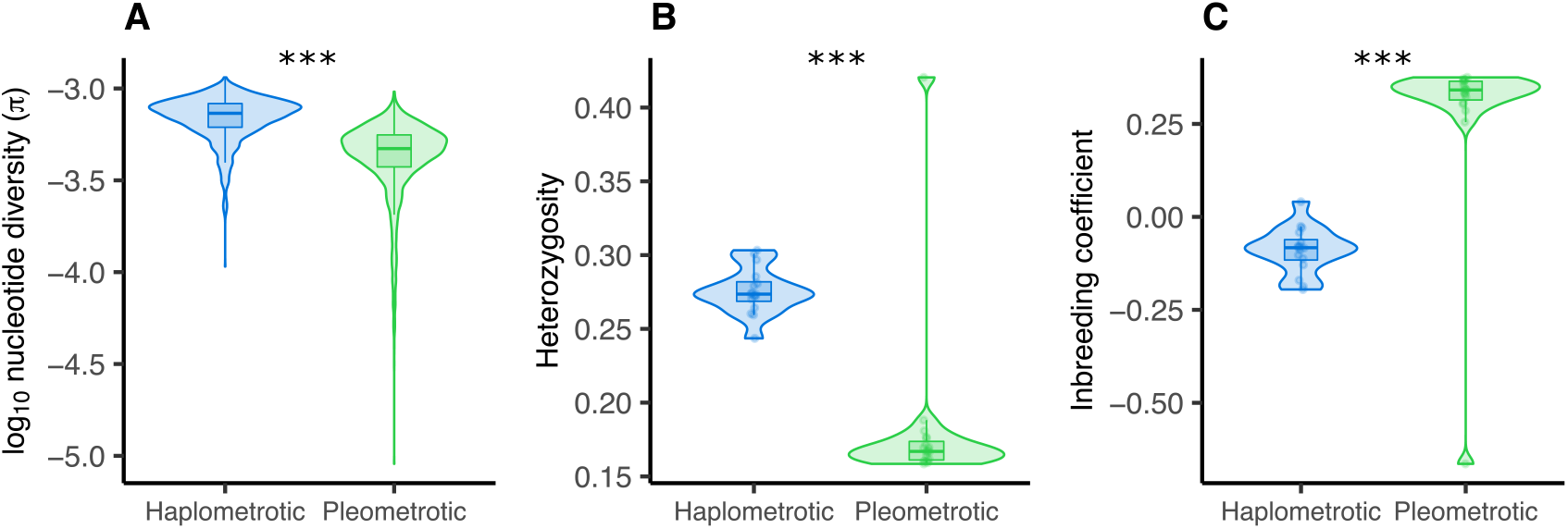
Higher Inbreeding and lower heterozygosity in the pleometrotic population. (A) Nucleotide diversity (𝜋), (B) heterozygosity and (C) inbreeding coefficient in the haplometrotic (blue) and pleometrotic (green) populations. Heterozygosity and inbreeding coefficient were calculated on a per individual basis, while 𝜋 estimates were calculated from 100-kb non-overlapping windows across the genome and averaged for each population. ***: P < 0.0001.

= 7.33e-4, median het = 0.274) (Figures 2A and 2B; 𝜋: Wilcoxon test: W = 3497100, p < 2.2e-16; het: Wilcoxon test: W = 288, p = 4.507e-07). Accordingly, inbreeding was highly prevalent in the P-population (median = 0.341) but absent in the H-population (median = -0.083) (Figure 2C; Wilcoxon test: W = 16, p = 4. 507e-07). Such genome-wide reductions of diversity in the pleometrotic population could be explained by a population bottleneck (founder effect), concordant with an overall positive Tajima’s *D* in both populations (Figure S3). This pattern is consistent with the repeated colonization and extinction of ephemeral habitats by *P. californicus*.

### The social polymorphism in P. californicus shows characteristics of a polygenic trait involving a supergene

Genome scans uncovered two scaffolds (22 and 23) that showed a very distinct pattern from the rest of the genome. Both scaffolds had high levels of genetic differentiation between the two social morphs, reduced nucleotide diversity, negative Tajima’s *D*, as well as very high extended haplotype scores (*xpEHH)* in the pleometrotic population (Figure S4). Further analyses revealed that SNPs were in high linkage disequilibrium (LD), with little to no LD decay in these scaffolds (Figure S5).

Linkage analysis on 108 haploid male offspring from a single queen belonging to the monogynous H-population using 2,980 ddRAD markers (double-digest Restriction-site Associated DNA sequencing) with no missing data put all markers in 29 Linkage groups (data not shown). The non-recombining region was found in linkage group 14 (339 RAD markers, 255 cM over 26 Mb; Figure 3A) and included scaffolds 22 and 23 as well as three other small scaffolds. This ∼8 Mb region contains 69 mapped markers in complete linkage (Figure S6). Hence, this is a non-recombining region co-segregating with the P- and H-populations, spanning across five complete physical scaffolds (22, 23, 38, 42, 48), and parts of scaffolds 6 and 14 (Table S2). Visual inspection of 182 markers from both males and queens showed that distinct haplotypes extended along the entire region, indicating absence of recombination (Figure 3A) across at least ∼8 Mb, consistent with patterns described for the supergenes in *Solenopsis* and *Formica* ants of non-recombining regions of 12.7 and 11 Mb, respectively [15,17]. Dense population genomic data using 5,363 SNPs genotyped across all 35 queens further confirmed this finding (Figure S7).

**Figure 3.**
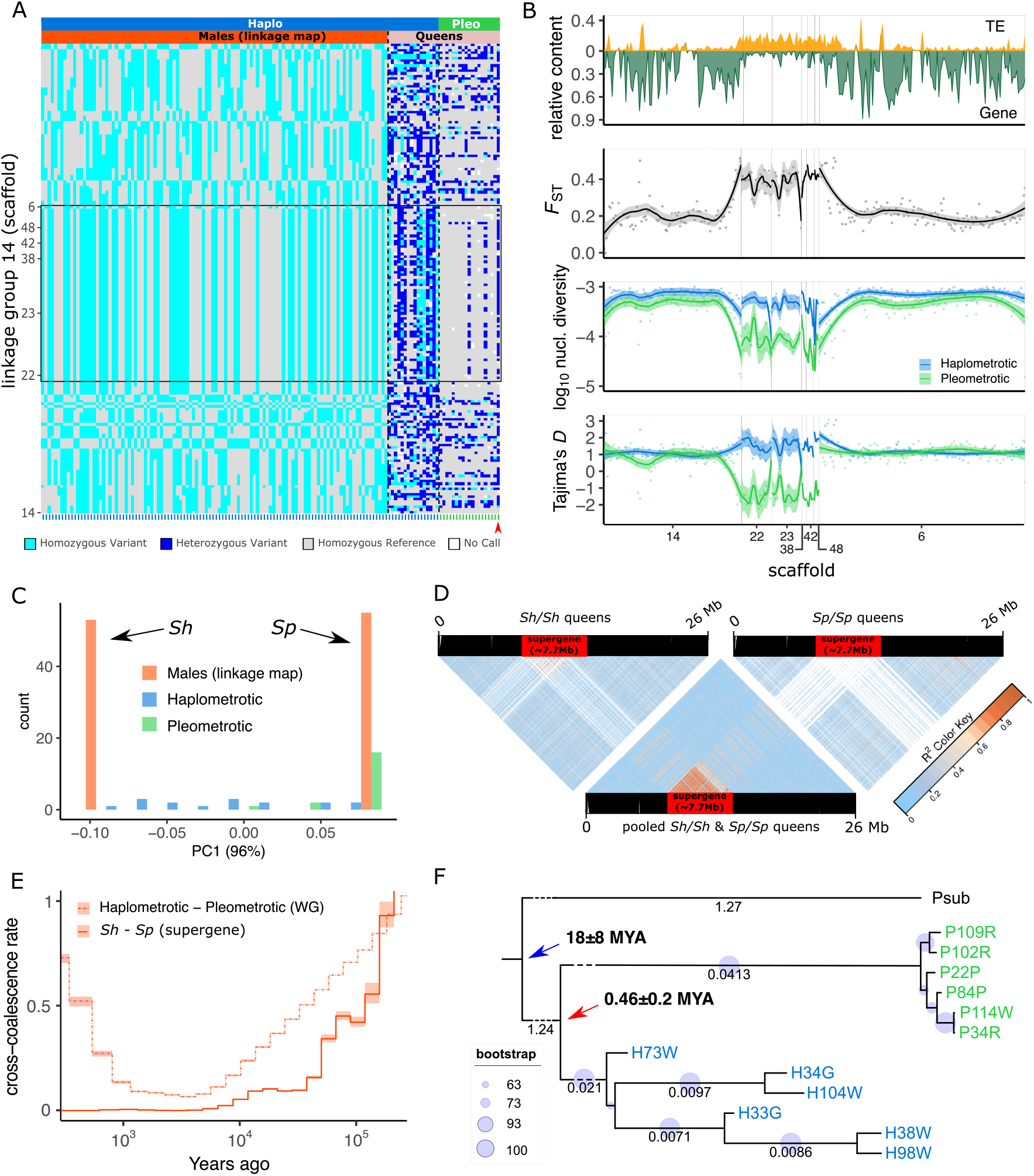
Genomic architecture and recombination rates at linkage group (LG) 14 harboring a putative supergene. (A) Genotypes for 108 sons from a single monogynous queen (haplometrotic population) based on RADseq markers and for the 35 queen samples from the pleometrotic and haplometrotic populations based on sites corresponding to RADseq markers from the whole genome resequencing data. Each row shows a SNP marker (182 in total) distributed across seven scaffolds on LG14. Columns (green ticks at the bottom for pleometrotic and blue ticks for haplometrotic) represent individual samples grouped by population (pleometrotic and haplometrotic) and dataset (males and queens). The black rectangle highlights the non-recombining putative supergene region, spanning scaffolds 22, 23, 38, 42, 48 and parts of scaffolds 6 and 14. Red arrowhead indicates the hybrid pleometrotic queen (P99P). (B) Sliding window analyses of LG14, showing (from top to bottom) the relative TE and gene content, genetic differentiation between populations (*F*_ST_), nucleotide diversity (𝜋) and Tajima’s *D* estimates in the pleometrotic (green) and haplometrotic (blue) populations. (C) Principal Component Analysis (PC1; 96%) based on 69 markers spanning the supergene region showing that the 108 male haplotypes are either Sh (negative PC1 scores) or Sp (positive PC1 scores). Diploid queens on the other hand are either homozygous for the Sp allele (most of pleometrotic queens, green bars) or heterozygous Sh/Sp (most of the haplometrotic queens, blue bars). (D) Linkage disequilibrium (LD) estimates (r^2^) across LG 14 for Sh homozygous queens (n=6), pooled queens homozygous for Sh (n=6) and Sp (n=6), and Sp homozygous queens only (n=6). The red rectangle shows the region of no recombination between the Sh and Sp haplotypes. SNPs are ordered according to physical position on the chromosome after lift over. Note that invariant marker sites are displayed in white, dominating particularly the homozygous Sh/Sh and Sp/Sp LD plots. (E) Demographic history of the supergene computed using MSMC2 with queens homozygous for the Sh (n=4) and Sp (n=4) alleles. The solid line shows the relative cross-coalescence rate (rCCR) at the supergene, while the dash-dotted line gives the genome-wide rCCR estimates. Lines and shaded areas are means and 95% confidence intervals, respectively. (F) Maximum-likelihood (ML) tree for the supergene region using 33,424 SNPs and *P. subnitidus* as outgroup species. Small branch length values at the tips of the tree were omitted for ease of visualization (Newick tree can be found in Dataset S6). The blue arrow indicates the upper bond estimate for the speciation of *P. subnitidus* and *P. californicus* 18 MYA (95% HPD ±8). The red arrow indicates the age estimate for the formation of the Sh–Sp haplotype groups 0.46 (95% HPD ±0.2) MYA.

Similar to the pattern described for the supergenes in *Solenopsis* and *Formica*, transposable element (TE) content in the *P. californicus* non-recombining region was significantly increased (Wilcoxon test: W = 14448, p < 2.2e-16), whereas exon content was significantly reduced compared to the rest of the same linkage group (Figures 3B and S8A; Wilcoxon test: W = 1602.5, p < 2.2e-16). This is consistent with the prediction that TEs accumulate in regions of low recombination [38,39]. Further, genetic differentiation as measured by *F*_ST_ between the H- and P-population was significantly increased in the supergene compared to the rest of the linkage group (Figures 3B and S8B; Wilcoxon test: W = 12261, p < 2.2e-16). Nucleotide diversity and Tajima’s *D* in this region were significantly reduced in the pleometrotic population (Figures 3B, S8C and S8D; Kruskal– Wallis rank sum test _nucleotide diversity_, 𝜒^2^ = 254.77, df = 3, p < 2.2e-16; Kruskal–Wallis rank sum test _Tajima’s *D*_, 𝜒^2^= 177.81, df = 3, p < 2.2e-16; results of pairwise Wilcoxon rank sum *post hoc* tests are displayed in the figure). Finally, 156 expressed transcripts from a previously published RNA-seq dataset [40] mapped to the supergene. 13 of those transcripts (8.3 %) were reported to be significantly associated with aggression depending on the social context of the queens (no significant enrichment, Dataset S2).

PCA on 69 markers along this region showed that PC1, which explains 96% of the variation, separates the male dataset of our mapping population into two distinct clusters, which we labeled as the Sh and Sp haplotype groups, indicating that these males are indeed the progeny of a heterozygous Sh/Sp mother queen (Figures 3C and S9). Superimposing SNP data from H- and P-queens to the PCA revealed that all but two pleometrotic queens (besides the hybrid queen) were homozygous for the Sp allele, while most haplometrotic queens were either heterozygous (Sh/Sp) or homozygous for the Sh allele (Figures 3C and S9). Further support for the suppression of recombination between Sh and Sp alleles came from LD analyses across queens. The region showed stronger LD in a mixed pool of Sh/Sh and Sp/Sp queens, as well as heterozygous Sh/Sp, in comparison to separate pools of Sh/Sh or Sp/Sp queens (Figures 3D and S10), indicating recombination in homozygous but not in heterozygous queens. The observed LD in homozygous Sh and Sp queens may be attributed to genomic rearrangements similar to *Formica* [16]. Suppression of recombination in heterozygotes is a key feature of supergenes [9,15,19,41]. Taken together, these data indicate the presence of an ∼8 Mb non-recombining supergene with two major haplotypes (Sh and Sp) segregating with the two populations of *P. californicus*. Unlike other social supergene systems (e.g. in *Solenopsis*) [7], there were no deviations from Hardy-Weinberg equilibrium (HWE) regarding the frequency of Sh/Sh, Sh/Sp and Sp/Sp genotypes at the supergene locus in either population (pleometrotic: 𝜒^2^(df = 1, N = 19) = 0.1396, p = 0.713; haplometrotic: 𝜒^2^ (df = 1, N = 16) = 0. 0.0711, p = 0.789) (Table S3).

Coalescence modelling revealed that the demographic histories of the supergene and the rest of the genome only diverged within the last 5,000 years. Then, cross-coalescence quickly increased across the genome, consistent with a secondary onset of gene flow between H- and P-populations, while remaining nearly zero in the supergene (Figure 3E). These findings suggest that Sh and Sp haplotypes initially diverged during the isolation period of the H- and P-population between approximately 200,000 and 5,000 years ago. During this period, one or multiple mutations suppressing recombination between Sh and Sp likely occurred (e.g. inversions), rendering the two haplotypes isolated, despite the admixture between the two populations. Consistent with a very recent origin of the supergene, phylogenetic analysis of Sh and Sp alleles with *P. subnitidus* as outgroup showed that Sh and Sp diverged less than 0.46 (95% highest probability density (HPD) 0.26-0.66) MYA (Figure 3F).

In addition to the supergene region, our genome scans revealed several regions under divergent selection between H- and P-populations. We first identified highly differentiated (outlier) regions by estimating *F*_ST_ in 100 kb non-overlapping windows across the genome (Figures S4 and S11) and subsequently compared estimates of Tajima’s *D* and nucleotide diversity in the two populations to identify signatures of selective sweeps (Figure S4). After clustering adjacent windows, we retained 51 clusters with negative Tajima’s *D* and reduced nucleotide diversity consistent with signatures of selective sweeps, of which 49 showed signatures of positive selection in the P-population and two in the H-population (Dataset S3). Cross-population extended haplotype homozygosity (*xpEHH*) tests [42] (Figure S4) identified 340 outlier SNPs in 32 genomic locations under selection (Dataset S4). Overlapping both analyses resulted in 17 remaining candidate regions under selection in the P-population (Table S4). Exonic SNPs in all 17 candidate regions included 70 synonymous changes and 73 non-synonymous variants affecting the protein sequence (Table 1).

**Table 1.** Effects of single nucleotide polymorphisms (SNPs) found in candidate regions that show strong signatures of selection in the pleometrotic population. We annotated 6,077 putative effects on protein coding genes of 3,347 SNPs.

| Effect | Count | Proportion [%] |
| --- | --- | --- |
| 3'UTR | 13 | 0.21 |
| 5'UTR | 1 | 0.02 |
| Downstream | 1391 | 22.89 |
| Intergenic | 3022 | 49.73 |
| Intron | 181 | 2.98 |
| Nonsynonymous variant | 73 | 1.20 |
| Synonymous variant | 70 | 1.15 |
| Upstream | 1319 | 21.70 |
| Other | 7 | 0.12 |

Twelve of these 17 regions (on scaffolds 14, 22 and 23) were in the supergene, while the remaining five regions with clear signatures of selective sweeps in the P-population were clustered on scaffolds 1, 9, 15, and 16 (Figure S4). These regions could initially not be placed into a linkage group due to the lack of ddRAD markers. To exclude regions potentially in linkage with the supergene, we constructed a relaxed linkage map with 11,986 (instead of 2,980) leniently filtered ddRAD markers (MAF ≥ 0.4 and 30% missingness), In this map, one of the candidate regions (scaffold 9) was contained in the supergene. The four remaining regions on scaffolds 1, 15 and 16 (Figure 4A), were placed into three different linkage groups, indicating that these regions are independent of the supergene.

**Figure 4.**
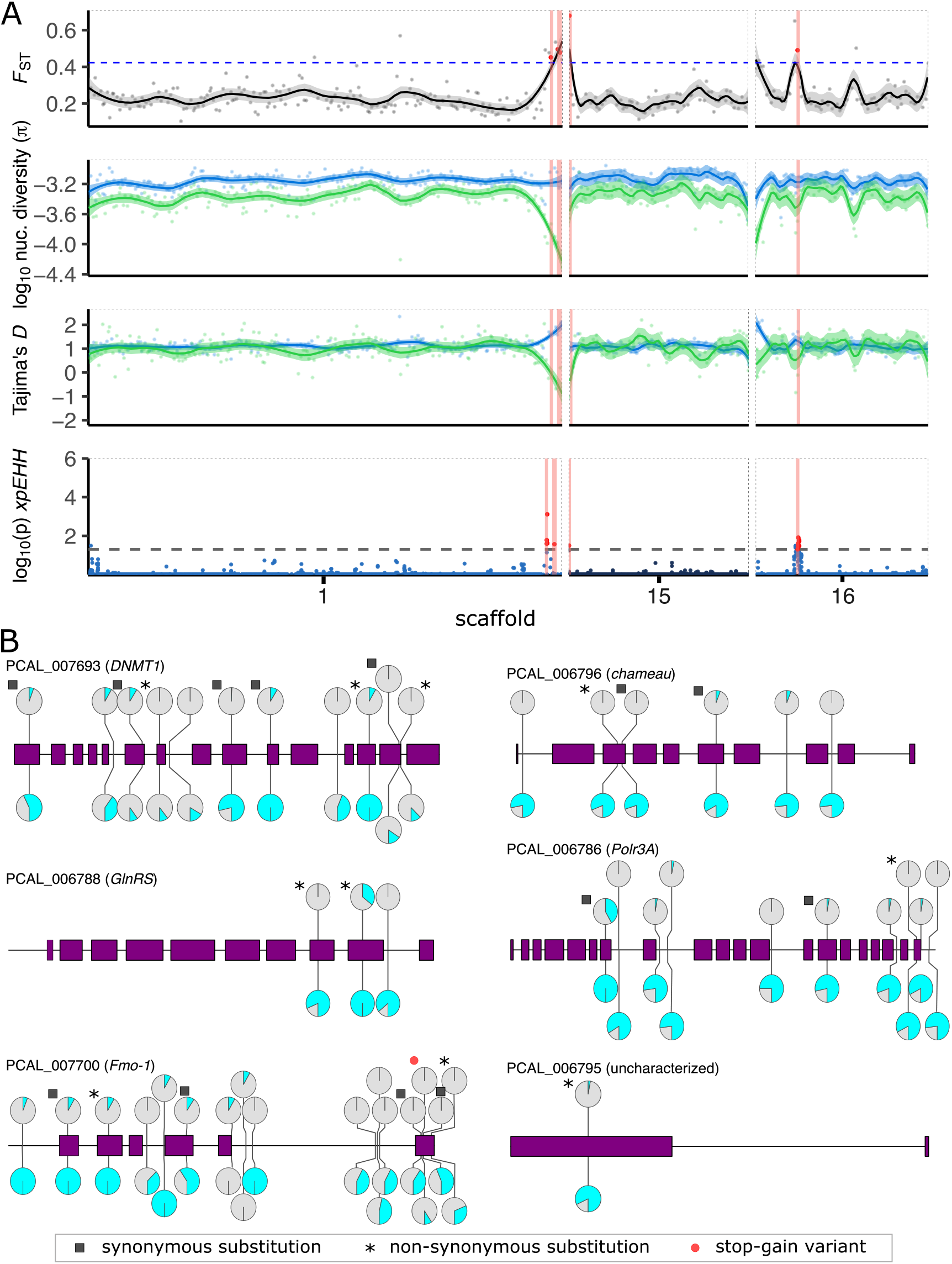
**Signatures of selection in non-supergene regions**. (**A**) Sliding window analyses of scaffolds 1, 15 and 16 showing (from top to bottom) genetic differentiation between populations (*F*_ST_), nucleotide diversity (𝜋), Tajima’s *D* and the cross-population extended haplotype homozygosity (*xpEHH*). Horizontal blue dashed line indicates *F*_ST_ threshold (*F*_ST_ > 0.423; red dots), while horizontal black dashed line represents threshold of significance for outlier SNPs (p < 0.05; red dots) in the *xpEHH* analysis. Four candidate regions in three scaffolds outside the supergene showing signatures of selection in the pleometrotic populations are highlighted in red in all panels. These regions are highly differentiated, have reduced diversity and Tajima’s *D* in the pleometrotic population and show a significant *xpEHH* peak. Lines and shaded areas are means and 95% confidence intervals, respectively. Data for other scaffolds (1-25) are shown in Figure S4. (**B**) Lollipop plots showing all variants affecting each of the six candidate genes (gene models in purple). The pie charts display frequencies of the reference (grey) and alternate (cyan) alleles in the pleometrotic (top; green background) and haplometrotic population (bottom; blue background).

Among the 48 genes in these four remaining regions, 34 are likely functional as they are not encoding TE proteins, are expressed, or homologous to other known insect proteins (Dataset S5). Six of these 34 genes are particularly promising candidates to sustain the social niche polymorphism, as they carry non-synonymous variants with a nearly perfect association of phenotype and genotype, in that >90% of haplometrotic individuals carry the alternative and >90% of pleometrotic individuals carry the reference allele (Figure 4B). Further, regions harboring the six genes exhibit *EHHS* scores revealing a pattern of increased extended haplotype homozygosity in the pleometrotic population (Figure S12), suggesting selective sweeps.

Visual inspection of variant positions showed that the most common pattern for all six genes is that pleometrotic queens are fixed for the reference allele (except for the hybrid P99P queen), while haplometrotic queens are either homozygous for alternate alleles or heterozygous (Figures S13A-S13F). This is consistent with a selective sweep fixing the reference allele in the P-population.

Apart from one uncharacterized protein (PCAL_006795) without functional annotation, the candidates encode evolutionarily well conserved proteins with functionally characterized orthologs in *D. melanogaster*. PCAL_006788 is orthologous to *GlnRS* (FBgn0027090), which encodes a tRNA synthetase involved in neurogenesis [43]. PCAL_006786 encodes an RNA polymerase III, orthologous to the transcriptional regulator Polr3A (FBgn0030687). PCAL_007700 encodes a flavin-containing monooxygenase homologous to *Drosophila* Fmo-1 (FBgn0034943) and Fmo-2 (FBgn0033079), which are suspected to be involved in xenobiotic detoxification and ageing [44]. The two remaining genes PCAL_006796 and PCAL_007693 particularly caught our attention, as they are orthologs of genes involved in epigenetic regulation of transcriptional activity. PCAL_006796 encodes chameau (ortholog to FBgn0028387), a MYST histone acetyl transferase involved in transcriptional regulation via histone modification [45]. PCAL_007693 encodes Dnmt1, a gene that is absent in Diptera. Dnmt1’s primary function is to maintain methylation patterns, and in social insects, inter-individual differences in DNA methylation have been linked to the epigenetic regulation of phenotypic plasticity, including caste determination in bees [46].

## Discussion

The social niche polymorphism [5] in colony founding of *P. californicus* was first reported over 20 years ago [28]. This polymorphism arises from differences in the social interaction among founding queens, which will eventually lead to fundamentally different colony structures, where either unrelated queens cooperate (i.e. primary polygyny) or a single queen monopolizes reproduction (i.e. primary monogyny) [11]. The principal change underlying the emergence of pleometrosis in *P. californicus* is hence the evolution of a lifelong tolerance for unrelated co-founding queens. Given that the majority of North American *Pogonomyrmex sensu stricto* are primarily monogynous, pleometrosis is likely a derived trait, adaptive under resource limitation and increased territorial conflicts [29]. Since its original description, we have learned a lot about behavioral, physiological, ecological, and socio-genetic correlates of haplo- and pleometrosis [11,26,27,29,32,33,47]. However, the genomic architecture and evolutionary history was unknown. The tight association of social morph and geographic location in *P. californicus* poses a unique challenge when exploring the genetic basis of this social polymorphism. As haplometrotic and pleometrotic individuals often occur in allopatry, any signature of genetic differentiation between both social morphs could be a product of genetic drift separating the populations. We therefore emphasize: While our findings are consistent with a young supergene underlying the social polymorphism in this species, our results are a characterization of genetic differentiation between two population that are almost fixed for distinct behavioral phenotypes. Removing this geographic confounder will require studying populations carrying both haplo- and pleometrotic individuals. However, despite our best efforts during several field seasons in the last years, we have not yet been able to find suitable populations. Thus, our study for now provides a first look at the genetic basis of social polymorphism in *P. californicus* under these limitations.

We identify an approximately 8 Mb non-recombining region, reminiscent of social supergenes described for social polymorphisms in *Solenopsis* fire ants [15] and *Formica* wood ants [16]. However, we also find genomic, population genetic, and ecological characteristics unique to the *Pogonomyrmex* system, highlighting the independent evolutionary histories of the social niche polymorphism in *Pogonomyrmex*, *Formica* and *Solenopsis*. The 8 Mb non-recombining region we identify is similar in size to the supergenes in *Formica* and *Solenopsis* (8 Mb vs. 11 and 12.7 Mb). According to estimates of *F*_ST_, genetic divergence between individuals from the two social forms is significantly higher within this region compared to other parts of the genome, which parallels the patterns reported for the other supergenes [48]. Tajima’s *D* and haplotype homozygosity statistics suggest a selective sweep of the Sp haplotype, although it could also be the result of a population bottleneck, similar to the *Solenopsis* Sb haplotype [49]. Very low LD decay and absence of recombinant F1 offspring of a heterozygous queen (Sh/Sp) suggest little to no recombination between Sh and Sp haplotypes at this locus. Further, TEs are significantly enriched at the supergene, as expected in the absence of recombination following a Muller’s ratchet dynamic [50].

Several other characteristics of the *P. californicus* social polymorphism differ from the well-studied examples in *Solenopsis* and *Formica* as well. In both, *S. invicta* and *F. selysi*, there is little evidence for genetic differentiation of polygyne and monogyne populations, with both forms often occurring in sympatry [51,52]. In *P. californicus* however, populations are fixed for either the haplo- or the pleometrotic strategy [11], with little gene flow between them. Accordingly, genetic differentiation between social forms in *P. californicus* is higher than in *S. invicta* and *F. selysi* [51,52], but it is similar to that observed in other *Formica* species, such as *F. exsecta* or *F. truncorum* [17]. The onset of gene flow observed between the haplometrotic and pleometrotic population starting ∼5,000 years ago could secondarily reduce genetic differentiation between both forms everywhere but in the non-recombining supergene, eventually resulting in the pattern also observed in *Solenopsis invicta* and *Formica selysi*, where genetic differentiation between both social morphs is restricted to the supergene [16,48].

We did not find significant deviations from HWE at the non-recombining region in either population, suggesting the absence of lethal haplotype combinations. This differs from *Formica* and *Solenopsis* where intrinsic (such as recessive lethal effects in *Solenopsis* and maternal killing effects in *Formica*) and extrinsic (such as green beard effects in *Solenopsis*) lethal combinations exist and are presumed crucial for the maintenance of the social niche polymorphism [53,54]. Further unlike in *Solenopsis* and *Formica*, we find additional genomic regions that segregate with the social polymorphism in *P. californicus*. Theoretical models for supergene evolution predict gradual establishment of supergenes by integration of formerly unlinked modifier loci [41,55]. According to these models, natural selection will favor genomic rearrangements that eventually join the supergene with previously unlinked but co-evolved loci carried on the same chromosome [56,57]. Hence, an alternative fate for the unlinked loci we describe here could be that they in fact persist as (e.g. population-specific) ‘modifier loci’, similar to the genetic architecture for refined Batesian mimicry described in *Papilio dardanus* [58–60] and *Heliconius numata* [61]. The presence of unlinked modifier loci influencing the expression and ameliorating negative fitness effects of major quantitative trait loci is not unusual [62], yet none have been described so far for the known social supergenes.

The association between the genotype at the non-recombining region and the social form is intriguingly imperfect in *P. californicus*. Pleometrotic queens were predominantly homozygous (85%) for a single haplotype (Sp), while most haplometrotic queens (93%) carried at least one copy of the Sh haplotype. This highlights that the genotype at this locus alone is insufficient to determine social form – a fundamental difference to the supergenes described in other ants [17,63]! The relatively young age of the suspected *Pogonomyrmex* supergene (*Pogonomyrmex*: ∼0.2 MYA, *Solenopsis*: 0.39–1.1 MYA [9,15,49] and *Formica*: 20–40 M years [17]) may contribute to this disparity. Alternatively, the regulation of social morphs in this species may involve a polygenic architecture, where the expression of haplometrosis or pleometrosis is influenced by social context, as well as environmental and ecological conditions [29]. This aligns with previous studies showing that environmental conditions and interactions with co-founding queens can modulate the expression of aggression and tolerance in different populations [32,33].

The four differentiated genomic regions outside the non-recombing region contain 34 genes that have undergone a selective sweep and potentially contribute to haplometrosis or pleometrosis. The nature of their interaction with the presumed supergene, whether additive or epistatic, remains unclear. Notably, the DNA methyltransferase *Dnmt1* and the histone acetyltransferase *chameau* exhibit a near-perfect correlation between genotype and phenotype, with all P-individuals carrying the reference allele and all H-individuals carrying the alternative allele. While these candidates suggest that epigenetic regulation modulates the social niche polymorphisms in *P. californicus*, in-depth functional studies will be necessary to explore how *chameau* or *Dnmt1* are indeed involved in the expression of haplo- and pleometrosis. As pleometrotic queens show a higher behavioral [32,33] and physiological [40] plasticity than haplometrotic queens during colony founding, differential epigenetic regulation of plastic traits like division of labor could maintain the two distinct social morphs.

## Conclusion

Pleometrosis and primary polygyny in *P. californicus* likely evolved as adaptation to challenging environments [29]. The genetic architecture associated with this social niche polymorphism shows similarities, but also significant differences compared to those described in e.g. *Formica* and *Solenopsis*. *Pogonomyrmex* also harbors a suspected non-recombining supergene, which is similar in size and has two major segregating haplotypes, as in the other systems. Additionally, four genomic regions showing signatures of a selective sweep in the pleometrotic population, potentially influence the expression of haplo- or pleometrosis. Further investigations are needed to fully understand the relevance and interactions of these loci. Unless both social morphs can be studied in sympatry, it will however remain challenging to conclusively resolve whether the suspected supergene and/or other genetic differences are indeed drivers of social polymorphism in this species or whether they have been shaped by demographic effects or local adaptations to some unknown ecological factor. Regardless, our study provides a first assessment of the genetic differentiation between haplo- and pleometrotic populations of *P. californicus*. It also showcases that the challenges with resolving the genetic basis for social niche polymorphisms, which may often be much more complex and diverse than previously appreciated.

## Materials and Methods

Further details about Materials and Methods are provided in the Supporting information file.

### Ant Collection

We collected gynes of *P. californicus* after their nuptial flights in 2018 and 2019 at Pine Valley (32.822 N, -116.528 W; pleometrotic population, P-population) and Lake Henshaw resort (33.232 N, -116.760 W; haplometrotic population, H-population), California, USA, which are 50 km apart.

### Genome sequencing

DNA libraries (NEBNext^®^ Ultra^™^ II FS DNA Libraries) of 16 queens from the H-population and 19 from the P-population were paired-end sequenced (2x 75 bp, average coverage 12x) at the Genomic Core Facility of the University of Münster (Germany), using a NextSeq 500 (Table S1).

### Genome assembly and annotation

We sequenced and assembled the genome of *P. californicus* (“Pcal2”) using data from MinION long reads (MinION, Oxford Nanopore Technologies, Oxford, UK), Illumina short reads, and previously published 10X Chromium linked reads [34]. For long read library preparation (Oxford Nanopore kit SQK-LSK109), we used 2 µg of DNA extracted from three white worker pupae from a pleometrotic colony (“Pcal18-02”). The library was sequenced for 48 h on a MinION, generating 1,650,688 reads that were trimmed and filtered with PoreChop (v.0.2.4) and FiltLong (v.0.2.0). We assembled the genome from ∼30x filtered Nanopore data using the following strategy. We created one assembly using CANU (v.2.0) [64] and a second assembly using wtdbg2 (v.2.5). Both assemblies were processed and optimized as follows: After contig assembly, we used SSPACE (v.3.0) [65] for scaffolding, LR_closer for gap filling, and ntedit for polishing with long-read data. We then used short-read, short-insert Illumina data from three samples (P50Y, P67P, P55R) for scaffolding with SSPACE before three rounds of polishing with Pilon [66]. The polished assembly was super-scaffolded with publicly available 10X Chromium data for *P. californicus* (accession SRX8051306) [34] using scaff10x and subsequent correction with break10x. Finally, we combined both assemblies with ntjoin, using the CANU-based assembly as reference and removed redundancies with funannotate clean (v.1.7.1).

We combined de-novo libraries from RepeatModeler2 (v.2.0.1) and EDTA (v.1.8.3) with public resources from RepBase (RepBase25.04.invrep.fa) and Dfam (Dfam3.0.arthropod.fa) for repeat annotation with RepeatMasker (v.4.0.7). Protein-coding and tRNA genes were annotated with funannotate v.1.7.1 in the soft-masked genome, including homology-based gene predictions from GeMoMa (v.1.7) [67] and supported by RNAseq data of 18 heads of *P. californicus* (Ernst et al., unpublished) and previously published long-read RNAseq data of a pool of workers [34].

### Read mapping and variant calling

Short-read data was cleaned with Trimmomatic (v.0.38) [68]. Paired reads were mapped to the genome assembly using BWA-MEM (v.0.7.17-r1188) [69]. After marking duplicates using Picard’s MarkDuplicates (v.2.20.0), we performed joint-variant calling across all samples with the GATK [70] and filtered variant calls with VCFtools (v.0.1.16) [71]. After a visual inspection of mapping quality of reads across the genome (Figure S14), we excluded fragmented scaffolds with low mapping quality and considered 613,411 SNPs identified in the 25 largest scaffolds (representing 88.2% of the assembly) for further analyses.

### Statistical phasing

We used SHAPEIT (v.2.r904) [72] to statistically phase the 25 largest scaffolds of the *P. californicus* genome based on 210,013 SNPs with no missing data (i.e. genotyped in all 35 queens), as no reference panel is available for *P. californicus*. Scaffolds 22 and 23 displayed an atypical architecture potentially representing a supergene (discussed further below) and were thus excluded (along with three other small scaffolds) for fineSTRUCTURE and MSMC2 analyses.

### Population structure analyses

LD-pruning with PLINK yielded 314,756 SNPs that we used for Principal Component Analyses (PCA), considering four eigenvectors. LD-pruned SNPs were further used for ADMIXTURE (v.1.3.0) analyses [73] with 2 to 4 clusters. To better characterize the relationships among queens in our sample, we used the haplotype-based approach implemented in fineSTRUCTURE [74]. In our model, we assumed a uniform recombination rate, based on the estimate of 14 cM/Mb found in *Pogonomyrmex rugosus* [75].

### Demographic population history analysis

We inferred demographic population history and the relative cross-coalescence rate (rCCR) using Multiple Sequential Markovian Coalescence modeling (MSMC2) [37] on four queens (i.e. eight phased haplotypes) from the haplometrotic (H123G, H118W, H73W and H123W) and pleometrotic populations (P22P, P134P, P117P and P109R). Mappability masks were generated using SNPABLE [http://lh3lh3.users.sourceforge.net/snpable.shtml]. We ran 1,000 bootstrap replicates, each time randomly sampling 60 Mb for each of the analyzed genomes. To obtain the time in years, we assumed a generation time of 4 years and a similar mutation rate as in honey bees [76], i.e. 3.6

* 10^-9^ mutations * nucleotide^-1^ * generation^-1^.

### Population genomic estimates

We used VCFtools (v.0.1.16) [71] to estimate nucleotide diversity (𝜋) (*--window-pi)* and Tajima’s *D (--TajimaD*) within each population using 100 kb non-overlapping windows across the genome. VCFtools’ *--het* function was employed to calculate heterozygosity and inbreeding coefficient on a per individual basis. Linkage disequilibrium (LD) decay was estimated by calculating pairwise r^2^ values between all markers within 200 kb windows using VCFtools’ *--hap-r2* function in each of the two populations. We used SNP markers with a minor allele frequency (MAF) of 0.2 to minimize the effects of rare variants and averaged r^2^ values between all pairs of markers in 100 bp distance bins for visualization.

### Genome scan for selection

We screened genomes of both populations for evidence of selection using an *F*_ST_ outlier approach [77–80] and by identifying regions of extended haplotype homozygosity (selective sweeps) [42,81], comparing the haplometrotic and pleometrotic populations. We excluded a total of six queens from pairs with relatedness > 0.4 and a potential F1 hybrid (P99P) from the analyses. We used mean *F*_ST_, nucleotide diversity *π*, and Tajima’s *D* estimated in 100 kb non-overlapping windows across the genome and from the empirical distribution of *F*_ST_ values, we considered the top 5% of windows as differentiated.

*F*_ST_ estimates can be affected due to variation in recombination rate across the genome [82]. Hence, we further applied the cross-population extended haplotype homozygosity (*xpEHH*) and the site-specific extended haplotype homozygosity (*EHHS*) tests [42,81], to detect signatures of selective sweeps. The *xpEHH* analyses were performed on the phased data from unrelated queens using the R package rehh (v.3.1.2) [83]. We used the Benjamani & Hochberg method for FDR correction and set a p-value of 0.05 as a threshold to identify outlier SNPs. Outliers occurring within 100 kb of each other were grouped, resulting in 32 clusters distributed across multiple scaffolds (Dataset S4). Regions within -/+ 100 kb around the most significant SNP of each cluster were regarded as candidates for positive selective sweeps. Regions identified by *F*_ST_ and by *xpEHH* overlapped at 17 genomic loci (Table S4).

### Characterization of candidate genes and SNPs

For functional characterization of genes, we performed BLAST (blastx) [84] searches against the nr database (January 2020), used OrthoFinder (v.2.3.1) [85] to identify *Drosophila melanogaster* orthologs, and used RepeatProteinMask to identify genes encoding TE proteins. We used SnpEff (v.5.0) [86] to annotate putative effects on protein-coding genes of all 3,347 SNPs contained in the candidate regions (Table 1). For the manual review of genes contained in genomic regions showing evidence for a selective sweep, we removed TE-encoded genes as well as genes without RNAseq support and no known homology to any proteins in the nr database, resulting in 34 putatively functional genes. Of these, 18 genes showed non-synonymous variants in our dataset (Dataset S5). In six of these 18 genes at least one non-synonymous variant showed a pattern where over 90% of the individuals from the H-population carried the alternative allele (i.e. either 0/1 or 1/1) and over 90% of the individuals from the P-population carried the reference allele (i.e. either 0/1 or 0/0) (Figure 4B).

### Haplotype frequencies

We calculated haplotype frequencies for the putative supergene region and tested for potential departure from Hardy-Weinberg equilibrium using a 𝜒^2^ test (Table S3). Haplotypes were assigned manually, based on visual inspection of the suggested supergene region (Figure S7); predominantly homozygous for the reference alleles (i.e. from a pleometrotic population) and alternate allele were assigned “Sp/Sp” and “Sh/Sh”, respectively, and predominantly heterozygous regions were assigned “Sh/Sp”.

### RAD sequencing and analysis

DNA of males collected from a monogynous colony at Lake Henshaw (H-population) in 2019 was used to construct reduced representation genomic libraries following Inbar et al. [87], using a modified double-digest (EcoR1 and Mse1) Restriction-site Associated DNA sequencing (ddRADseq) protocol [88,89]. Briefly, genomic DNA was digested at 37°C for 8hrs and ligated with uniquely barcoded adaptors. The adaptor ligated products were PCR amplified, pooled, and size-selected using Beckman Coulter, Agencourt AMPure XP beads. Libraries were sequenced in one lane of a paired-end, 150 bp reads, on a HiSeq X Ten Illumina sequencer.

We used fastQC (v.0.11.7), Trimmomatic (v.0.39) [68], and *process_radtags* (STACKS v.2.2) [90] for quality control, filtering, and demultiplexing the raw data. Reads were mapped with Bowtie2 v2.3.4.2 (Langmead et al 2012) and alignments processed with *ref_map.pl* and *populations* (STACKS v.2.2), resulting in 55,340 raw variants. After filtering with VCFtools (v.0.1.16) [71] and excluding heterozygous markers we arrived at a final set of 2,980 markers used for building a genetic map. Markers were mapped with MultiPoint [http://www.multiqtl.com] using the backcross population setting. A maximum threshold recombination frequency of 0.25 resulted in 29 linkage groups (data not shown). A skeleton map was generated using delegate markers and Kosambi’s mapping function was used to convert recombination frequencies to centimorgans [91].

### Characterization of the supergene

To investigate the potential supergene identified in the current study, we used SNP markers from queens (whole genome resequencing) and males (RADseq). We used BCFtools’ (v.1.9) *isec* and *merge* functions to extract 182 shared SNPs between the two datasets found on LG14. Genotypes of the males and queens at LG14 were visualized using VIVA (v.0.4.0) [92]. To further characterize the two alleles of the supergene, we performed a PCA on 69 SNP markers identified in the non-recombining area of LG14 (i.e. scaffolds 22, 23, 38, 42, 48, and parts of 6 and 14) in datasets of males and queens.

To investigate recombination suppression in the supergene region, we used SNP data from six queens homozygous for the Sh allele (H104W, H33G, H34G, H38W, H73W and H98W) and six others homozygous for the Sp allele (P22P, P102R, P114W, P109R, P34R, P84P), assigned based on visual inspection of their genotypes (Figure S7). After VCF lift over (on scaffolds 14, 22, 23, 38, 42, 48 and 6 into LG14) using flo [48] and picard’s *LiftoverVcf* (v.2.20.0), we used VCFtools (v.0.1.16) [71] for filtering and LDheatmap (v.1.0-4) [93] to visualize pairwise linkage disequilibrium (LD). We calculated gene and TE content in 100 kb sliding windows across LG14 using our *P. californicus* gene and TE annotations, respectively. To explore whether any of the transcripts previously shown to be associated with the aggressive behavior of queens [40] belong to the supergene, we aligned the assembled transcripts to the Pcal2 reference genomes using GMAP [94]. We only retained the top hits with at least 70% query coverage, yielding 7303 (out of 7890) transcripts unambiguously aligned to the reference genome. We then used BEDtools (v.2.29.2) [95] to extract transcripts that mapped to the supergene region (Dataset S2).

### Dating the supergene formation

We estimated the divergence between Sh and Sp by computing the rCCR, using MSMC2 [37] on four Sh/Sh queens (H104W, H38W, H73W and H98W) and four Sp/Sp queens (P114W, P50R, P84P and P97R). We limited the analysis to the supergene region (scaffolds 22, 23 and part of scaffold 6) and ran 500 bootstraps by randomly sampling 2 Mb each time.

Further, we performed a phylogenetic analysis of Sh and Sp alleles with *P. subnitidus* as an outgroup species. Sequencing data of one *P. subnitidus* individual was generated and processed as described above for *P. californicus*. Joint SNP calling was performed using GATK [70] and a maximum-likelihood (ML) tree was constructed based on 33,424 SNPs at the supergene region (scaffolds 22, 23, 38, 42, 48 and parts of scaffolds 6 and 14), using RAxML [96] under the GTRGAMMA model with bootstrapping.

We then used a fossil-calibrated dated phylogeny [97] to date the split between the Sh and Sp haplotype groups using a phylogenetic approach. The Ward et al. [97] phylogeny dated the split between *P. vermiculatus* and *P. imberbiculus* at 18 (95% HPD ±8) MYA. This is an upper bound on the speciation date of *P, californicus* and *P. subnitidus* according to the most recent *Pogonomyrmex* phylogeny [98]. We calculated a ratio of 1:39 between the speciation time of *P. subnitidus* and *P. californicus*, and the Sh–Sb haplotypes divergence time. Based on our dating reference (i.e. 18±8 MYA), we obtained 0.46 (95% HPD ±0.20) MYA as an upper bound for the divergence between the two haplotype groups, which is consistent with the MSMC2 analysis. We note that the confidence in the age inference is strengthened by the agreement between two distinct approaches: the inference by MSMC2 is based on a coalescent model, which is a distinct approach from the phylogenetic dating approach. The former approach depends on estimates of mutation rate and generation time, whereas the latter depends on fossil calibration points. The tree figure was produced using interactive Tree Of Life (iTOL) [99].

## Data Availability

The genome assembly and the raw sequencing data prior to trimming and mapping are available at NCBI (BioProject: PRJNA682388; release pending acceptance). A detailed description of our bioinformatic analysis pipelines is publicly available at https://github.com/merrbii/Pcal-SocialPolymorphism.

## Acknowledgments

We thank Pine Valley Academy (Pine Valley, California, USA) for support and the staff of Lake Henshaw resort (Lake Henshaw, California, USA) for permission to collect ants on their premises. Thanks to Jenny Märzhäuser, Tobias van Elst and Ti Eriksson for help in collecting the ants, and to Jennifer Fewell for hosting us at Arizona State University, Tempe, Arizona, USA. We are grateful for Hilde Schwitte’s and Una Hadziomerovic’s help in DNA extraction, barcoding, and library construction. JG and UE were funded by the German Research Foundation (DFG) as part of the SFB TRR 212 (NC³) – TP C04 project numbers 316099922 and 396780988. ME was funded by the German Research Foundation (DFG) – 403813881 with grant to LS (SCHR 1554/2-1) under the priority program “Rapid evolutionary adaptation: Potential and constraints” (SPP 1819). ME acknowledges support by the DFG Research Training Group 2220 “Evolutionary Processes in Adaptation and Disease”.

## Author Contributions

JG, UE, ME and LS designed the research, UE organized sample collection and behavioral assays, AL, LS, ME, UE, EP and JG analyzed data, ME, UE, LS and JG wrote the manuscript, JG and LS coordinated the project, all authors revised the manuscript.

