## Supporting information file for "Evolutionary genomics of socially polymorphic populations of *Pogonomyrmex californicus*"

[‡]: equal contribution

**This Supporting information includes:**

Extended Material and Methods
Figures S1 to S15
Tables S1 to S4
Legends for Datasets S1 to S6
Supplemental References

**Material and Methods**

*Ant Collection*

We collected gynes of *P. californicus* after their nuptial flights in 2018 and 2019 at Pine Valley (32.822 N, -116.528 W; pleometrotic population, P-population) and Lake Henshaw resort (33.232 N, -116.760 W; haplometrotic population, H-population), California, USA, which are 50 km apart. Queens were snap-frozen in liquid nitrogen and transferred to vials containing 0.5 ml RNAlater (Invitrogen). There are no legal regulations concerning the collection or treatment of ants in the USA or Germany. All ants were barcoded using *Cytochrome Oxidase I* (COI) to confirm species identification and determine mitochondrial variation in both populations.

*DNA extraction, library prep and genome sequencing*

DNA was extracted using phenol-chloroform following [1]. Libraries were constructed using NEBNext® Ultra™ II FS DNA Library Prep Kit for Illumina (NEB #E7805; New England BioLabs, Ipswich, USA) according to manufacturer's protocol. DNA extracts and libraries were controlled for purity and size using Nanodrop and Bioanalyzer 2100 (Agilent, Santa Clara, USA). Libraries were paired-end sequenced (2x 75 bp) at the Genomic Core Facility of the University of Münster (Germany), using a NextSeq 500 system with NextSeq 500 High Output Kit v2.5 (Illumina, San Diego, USA). We sequenced 16 queens from the H-population and 19 from the P-population at an average coverage of 12x (Supplemental file, Table S1).

*Genome assembly and annotation*

We sequenced and assembled the genome of *P. californicus* ("Pcal2") using data from MinION long reads (MinION, Oxford Nanopore Technologies, Oxford, UK), Illumina short reads, and previously published 10X Chromium linked reads [2]. For long read library preparation, we sampled three white worker pupae from a pleometrotic colony ("Pcal18-02") in liquid nitrogen for high-molecular weight DNA extraction using a phase-separation protocol, optimized for DNA extraction from ants. After quality control with agarose gels, Nanodrop and Qubit measurements, we used 2 µg of DNA for library preparation with the Ligation Sequencing Kit (Oxford Nanopore) SQK-LSK109.

The resulting library was sequenced for 48 h on a MinION sequencer, using a single flow cell. After basecalling with guppy (v.4.0.14) we retrieved 1,650,688 reads (mean read length 4.9 kb) with a mean read quality of 13.4. Read length N50 was 18.5 kb and a total of 8.1 Gb of data was generated. We used PoreChop (v.0.2.4) to remove adapter contaminations from MinION reads before further filtering the data with FiltLong (v.0.2.0), using paired-end Illumina data from three samples (P50Y, P67P, P55R). We assembled the genome from ~30x filtered Nanopore data using the following strategy. We created one assembly using CANU (v.2.0) [3] and a second assembly using wtdbg2 (v.2.5). Both assemblies were processed and optimized as follows: After contig assembly, we used SSPACE (v.3.0) [4] for scaffolding, LR_closer for gap filling, and ntedit for polishing with long-read data. We then used short-read, short-insert Illumina data from three samples (P50Y, P67P, P55R) for scaffolding with SSPACE before three rounds of polishing with Pilon. The polished assembly was super-scaffolded with publicly available 10X Chromium data for *P. californicus* (accession SRX8051306, Bohn et al. 2021) using scaff10x and subsequent correction with break10x. We used an in-house, blastn-based pipeline to remove bacterial contaminations from each assembly. Finally, we combined the CANU-based and the wtdbg-based assemblies with ntjoin, using the CANU-based assembly as reference. The joint assembly was filtered for redundancy using funannotate clean (v.1.7.1) (https://github.com/nextgenusfs/funannotate/tree/1.7.1), removing four scaffolds.

*Genome annotation*

We annotated repeats and transposable elements (TEs) in the genome using a combination of de-novo, structural and homology-based approaches. We ran RepeatModeler2 (v.2.0.1) and EDTA (v.1.8.3) on the genome assembly to generate de-novo repeat libraries. Libraries from both tools were combined and redundancies removed with CD-HIT (v.4.8.1). We classified repeats using a combination of PasteClassifier (included in REPET v.2.5), DeepTE (github.com/LiLabAtVT/DeepTE) and TEsorter (v.1.2.5.2) (https://github.com/zhang renga ng/TEsorter). We removed putative host proteins from the library by blasting all unclassified repeats against uniprot_arthropoda (uniprot-arthropoda+6656 downloaded 27-Apr-2018) with “blastx -

soft_masking false -evalue 1e-10". We combined our *de novo* library with public resources from RepBase (RepBase25.04.invrep.fa) and Dfam (Dfam3.0.arthropod.fa). Finally, using this combined library, we annotated the genome with RepeatMasker (v.open-4.0.7) with -s -gff -a -inv -small -excln -cutoff 250 -nolow. The produced annotation was subsequently improved further with One-code-to-find-them-all (v.1.0) [5].

Protein-coding and tRNA genes were annotated with funannotate (v.1.7.1) in the soft-masked genome. We used short-read Illumina paired-end RNAseq data of 18 heads of *P. californicus* (Ernst et al., unpublished) and previously published long-read minION RNAseq data of a pool of workers [2] for training. Homology-based gene predictions were included by running GeMoMa (v.1.7) [6] on a set of five published ant genomes (*Pogonomyrmex barbatus*, GCF_000187915.1; *Ooceraea biroi*, GCF_003672135.1; *Camponotus floridanus*, GCF_003227725; *Solenopsis invicta*, GCF_000188075.2, *Acromyrmex echinatior*, GCF_000204515.1). funannotate initially produced 26,408 gene models. After functional annotation with interproscan [7], EggNog [8], and funannotate, we removed any protein-coding gene that did not have any functional annotation or known ortholog and did not have any RNAseq coverage. Of the predicted genes, 9,435 had at least some RNAseq support and 15,110 showed at least some homology to known proteins (with E-value < 1E-5 against UniRef90 Release: 2020_04, 12-Aug-2020). 1,279 genes had RNAseq support but were not homologous to any protein in UniRef90.

<u>*Read mapping and variant calling*</u>

We used FastQC (v.0.11.7) to inspect the quality of the raw short read data. Short and low quality reads were removed using Trimmomatic (v.0.38) (parameters: *SLIDINGWINDOW*:4:20 *MINLEN*:40) to only keep paired and high quality reads [9]. The resulting paired reads were mapped to the newly produced assembly of *P. californicus* (Pcal2) using BWA-MEM (v.0.7.17-r1188) with default parameters [10]. We assessed the quality of the resulting alignments using QualiMap (v.2.2.1) [11].

After marking duplicates using Picard's MarkDuplicates (v.2.20.0), we performed joint-variant calling across all samples, in two rounds.

We first generated a high quality set of SNPs that we used for base quality recalibration in the second pass. In each pass, we used GATK's HaplotypeCaller (v.4.1.2.0) with default parameters [12]. Using this approach, we recovered a total of 1,486,819 SNPs that we hard-filtered using the GATK VariantFiltration tool with the following parameters: the variant confidence normalized by depth (QD < 2.0), estimates of the variant strand bias test (FS > 60.0 and SOR > 3.0), mapping quality (MQRankSum < -5.0 and MQ < 40.0) and the rank sum test for the positioning of variants within (readsReadPosRankSum < -5.0). We based our choice of these parameters on a visual inspection of the distribution of annotation values of the raw SNPs as recommended by the GATK documentation [13].

The 1,359,080 SNPs that passed the applied hard filters were further filtered using VCFtools (v.0.1.16) [14] and we kept only biallelic SNPs occurring in at least 85% of samples, with a minor allele count (MAC) ≥ 3 and a site depth of 3X to 24X, resulting in 646,060 SNPs. These cut-offs were chosen based on the distribution of these parameters in the raw SNPs left after applying the hard filters. After a visual inspection of read mapping quality across the genome (Supplemental file, Figure S14), we excluded short scaffolds with low mapping quality and considered 613,411 SNPs identified in the 25 largest scaffolds (representing 222.6 Mb, i.e. 88.2% of the assembly) for further analyses.

<u>*Statistical phasing*</u>

We used SHAPEIT (v.2.r904) [15] to statistically phase the 25 largest scaffolds of the *P. californicus* genome based on our resequencing data of 35 single individuals, as no reference panel is available for *P. californicus*. To produce a reliable set of phased genomes, we only considered 210,013 SNPs with no missing data (i.e. genotyped in all 35 queens) for phasing. We separated the SNP dataset genotyped across all 35 individuals into scaffolds using BCFtools (v.1.9) and performed phasing by running SHAPEIT (*-check*, *-phase* and *-convert*) on each scaffold separately.

Scaffolds 22 and 23 displayed an atypical architecture potentially representing a supergene (discussed further in the manuscript). Therefore scaffolds 22 and 23 as well as three other small scaffolds were excluded, and we only used phased SNPs contained in the 20 largest scaffolds for fineSTRUCTURE and MSMC2 analyses.

*Population structure analyses*

We used PLINK to LD-prune the 613,411 SNPs at a threshold of $r^2 < 0.2$, using the *--indep-pairwise* 1 kb 1 0.2 command. This yielded 314,756 LD-pruned SNPs that we then used for Principal Component Analyses (PCA) using the *--pca* command of PLINK (v.1.90p) [16,17], considering four eigenvectors. The LD-pruned SNPs were further used for ADMIXTURE (v.1.3.0) analyses [18] with 2 to 4 clusters. The best fit was selected based on cross-validation error rates.

To better characterize the relationships among queens in our sample, we used the haplotype-based approach implemented in fineSTRUCTURE [19]. In this method, each individual is considered in turn as a recipient, whose chromosomes are reconstructed as a combination of haplotypes from all other individuals (the donors), using ChromoPainter [19]. Closely related individuals are expected to share more haplotype blocks with each other. The final output is a coancestry matrix that can be visualized as a heat map summarizing the number of shared haplotypes between all pairs of individuals in the analyzed data sets.

We first converted SHAPEIT output to ChromoPainter's PHASE format using the *impute2chromopainter.pl* provided by the developers of fineSTRUCTURE. We ran fineSTRUCTURE (v.4.1.0) under the 'linked' mode with default parameters. In our model, we assumed a uniform recombination rate, based on the estimate of 14 cM/Mb found in *Pogonomyrmex rugosus* [20]. The coancestry matrix plot was produced in R (v.4.0.2) [21] using an R script, also provided with fineSTRUCTURE [https://people.maths.bris.ac.uk/~madjl/finestructure/finestructureR.html].

*Demographic population history analysis*

We inferred demographic population history from individual genomes using Multiple Sequential Markovian Coalescence modeling (MSMC2) [22]. We used phased data from four single individual queens (i.e. eight phased haplotypes) from the haplometrotic (H123G, H118W, H73W and H123W) and pleometrotic populations (P22P, P134P, P117P and P109R).

Mappability masks for each scaffold were generated using SNPABLE [http://lh3lh3.users.sourceforge.net/snpable.shtml].

Effective population size ($N_e$) and the relative cross-coalescence rate (rCCR) were then calculated using MSMC2 following the developer's guidelines [22,23]. We ran 1,000 bootstrap replicates, each time randomly sampling 60 Mb for each of the analyzed genomes. To obtain the time in years, we assumed a generation time of 4 years and a similar mutation rate as in honey bees [24], i.e. 3.6 * $10^{-9}$ mutations * nucleotide$^{-1}$ * generation$^{-1}$.

*Population genomic estimates*

We used VCFtools (v.0.1.16) [14] to estimate nucleotide diversity ($\pi$) (*--window-pi)* and Tajima's *D* *(--TajimaD*) within each population using 100 kb non-overlapping windows across the genome. VCFtools' *--het* function was employed to calculate heterozygosity and inbreeding coefficient on a per individual basis. To reduce the effects of mapping errors, windows where the average mapping quality is below 40, and/or show extreme depth of coverage and/or SNP number were excluded from further analyses (Supplemental file, Figure S15).

Linkage disequilibrium (LD) decay was estimated by calculating pairwise $r^2$ values between all markers within 200 kb windows using VCFtools' *--hap-r2* function in each of the two populations. We used SNP markers with a minor allele frequency (MAF) of 0.2 to minimize the effects of rare variants. We next averaged $r^2$ values between all pairs of markers in 100 bp distance bins for ease of visualization.

We tested for statistical differences between populations by performing a Wilcoxon rank sum test with "Bonferroni" adjustments of the p-values in R (v.4.0.2) [21].

*Genome scan for selection*

To identify candidate genes potentially responsible for the social polymorphism in *P. californicus*, we screened genomes of both populations for evidence of selection using an $F_{ST}$ outlier approach and by identifying regions of extended haplotype homozygosity (selective sweeps), comparing the haplometrotic and pleometrotic populations [25–28].

Prior to computing $F_{ST}$ values between the two populations, we estimated relatedness between queens using PLINK's *--make-rel* function [16,17]. Based on visual inspection of the distribution of relatedness values, we excluded queens from pairs with relatedness > 0.4 leaving us with 17 (of 19) and twelve (of 16) queens from the pleometrotic and haplometrotic populations, respectively. Moreover, we excluded one queen from the pleometrotic population as our population structure analysis indicated that it was a F1 hybrid (P99P).

We calculated mean $F_{ST}$ in 100 kb non-overlapping windows across the genome using VCFtools' functions *--weir-fst-pop* and *--fst-window-size* (v.0.1.16) [14]. From the empirical distribution of $F_{ST}$ values, we considered the top 5% of these windows as significantly differentiated (Supplemental file, Figure S11).

To determine in which population the differentiated windows are under selection, we computed nucleotide diversity $\pi$ and Tajima's $D$ in 100 kb non-overlapping windows across the genome (Supplemental file, Figure S4) and reasoned that these regions should show a reduced Tajima's $D$ (typically negative values) and nucleotide diversity in the population where they are subject of selection. Genomic windows failing these conditions were flagged as ambiguous and were discarded from further processing, resulting in a total of 51 candidate regions evolving under divergent selection between the populations (Dataset S3).

$F_{ST}$ estimates can be affected due to variation in recombination rate across the genome [29]. Hence, we further applied the cross-population extended haplotype homozygosity (*xpEHH*) and the site-specific extended haplotype homozygosity (*EHHS*) tests [30,31], to detect signatures of selective sweeps. The *xpEHH* compares the length of haplotypes between populations to detect selective sweeps in which the selected allele has approached or reached fixation in at least one population [30]. The *EHHS* on the other hand is defined as the decay of identity of haplotypes surrounding a core marker of a population as a function of distance [31].

The *xpEHH* analyses were performed on the phased data from unrelated queens using the R package rehh (v.3.1.2) [32] with the following settings: *polarize_vcf=FALSE*, *min_maf=0.05*, *scalegap=20,000* and *maxgap=200,000*. The latter two options were used in order to reduce false positives caused by large gaps between markers [33]. We used the Benjamani & Hochberg method for FDR correction and set a p-value of 0.05 as a threshold to identify outlier SNPs. Outliers occurring within 100 kb of each other were clustered into one outlier region under selection, resulting in 32 clusters distributed across multiple scaffolds (Dataset S4). Regions that are within -/+ 100 kb around the most significant SNP of each cluster were kept as candidates for positive selective sweeps.

We used BEDtools' *intersect* function (v.2.29.2) [34] to extract regions that overlapped between the two selection screen approaches, producing 17 candidate regions (Supplemental file, Table S4). We excluded an ambiguous region on scaffold 11 (position: 1,200,000-1,349,402) that showed opposing pattern of selection (i.e. $F_{ST}$-based approach suggested selection in the pleometrotic population while the *xpEHH* indicated selection in the haplometrotic population).

*Characterization of candidate genes and SNPs*

We used BEDtools' *intersect* function (v.2.29.2) [34] to extract regions identified simultaneously by the $F_{ST}$- and *xpEHH*-based approach. Next, we used SnpEff (v.5.0) [35] to annotate putative effects on protein-coding genes of all 3,347 SNPs contained in the candidate regions (Table 1).

The $F_{ST}$- and *xpEHH*-based approaches to detect selection yielded a set of 145 candidate genes (including 97 in the supergene region). To identify putative function of these candidate genes, we performed a BLAST (blastx) [36] search against the nr database (January 2020). In addition to BLAST, we used OrthoFinder (v.2.3.1) [37] to identify *Drosophila melanogaster* orthologs to our candidate genes. Further, we used RepeatProteinMask to identify genes encoding TE proteins. For the manual review of the 48 genes contained in genomic regions showing evidence for a selective sweep outside the supergene, we removed TE-encoded genes as well as genes without RNAseq support and no known homology to any proteins in the nr database, resulting in 34 putatively functional genes (Dataset S5). Of these, 18 genes showed non-synonymous variants in our dataset. In six of these 18 genes at least one non-synonymous variant showed a pattern where over 90% of the individuals from the H-population carried the alternative allele (i.e. either 0/1 or 1/1) and over 90% of the individuals from the P-population carried the reference allele (i.e. either 0/1 or 0/0) (Figure 4B).

*Haplotype frequencies*

We calculated haplotype frequencies for the putative supergene region and tested for potential departure from Hardy-Weinberg equilibrium using a $\chi^2$ test (Supplemental file, Table S3). Haplotypes were assigned manually, based on visual inspection of the suggested supergene region (Supplemental file, Table S2 and Figure S5); predominantly homozygous for the reference alleles (i.e. from a pleometrotic population) and alternate allele were assigned "Sp/Sp" and "Sh/Sh", respectively, and predominantly heterozygous regions were assigned "Sh/Sp".

*RAD sequencing and analysis*

Winged males from a monogynous colony at Lake Henshaw (H-population) were collected during the mating season 2019 and immediately stored in 100% ethanol. DNA was isolated using a phenol-chloroform method [1]. Tests using six microsatellite markers confirmed that all males belonged to the same matriline (results not shown). Following Inbar et al. [38], we constructed a reduced representation genomic library using an adjusted double-digest Restriction-site Associated DNA sequencing (ddRADseq) protocol (Parchman et al. 2012, Peterson et al. 2012). First, we digested DNA withEcoR1 as rare-cutter and Mse1 as frequent-cutter restriction enzyme. Then, we ligated DNA fragments to barcoded adaptors for multiplexing (using CutSmart buffer NEB). The adaptor-ligated products were PCR amplified, pooled, and size-selected using Beckman Coulter, Agencourt AMPure XP beads. The libraries were sequenced in one lane of a paired-end, 150 bp reads, on a HiSeq X Ten Illumina sequencer.

Quality of a total of 369,727,466 paired-end RADseq reads was assessed with fastQC (v.0.11.7). We used Trimmomatic v.0.39 [9] to remove adaptor sequences with the following settings: *ILLUMINACLIP:TruSeq3-PE-2.fa:2:30:10:8:true SLIDINGWINDOW:4:15 MINLEN:100*, before running *process_radtags* (STACKS v.2.2) [41] for demultiplexing and further filtering.

Mapping to the genome assembly of *P. californicus* (Pcal2) was performed using BWA-MEM (v.0.7.17-r1188) with default parameters [10]. The resulting alignments were processed with *ref_map.pl* and *populations* (STACKS v.2.2) resulting in 55,340 raw variants. We used VCFtools (v.0.1.16) [14] to only keep biallelic markers without missing data (i.e. genotyped in all males) and a MAF ≥ 0.45, yielding 4,082 markers. Finally, we excluded heterozygous markers (likely sequencing errors as males are haploid) producing a final set of 2,980 markers used for building genetic maps.

The 2,980 markers were all mapped with MultiPoint [http://www.multiqtl.com] using the Backcross population setting. A maximum threshold rfs value of 0.25 resulted in 29 linkage groups. Note, MultiPoint implemented a feature that recognizes markers showing linkage in the opposite phase (to automatically recognize phasing errors in large data sets), hence it is no longer necessary to first double the markers, switch phase for one marker set and then start mapping as described in [42]. Multipoint linkage analysis of loci within each linkage group (LG) refined marker order through resampling (jack-knifing, [43]). A skeleton map was generated using delegate markers for all markers that showed 100% linkage. Kosambi's mapping function was used to convert recombination frequencies to centimorgan [44]. The complete map will be published elsewhere, for this manuscript we are only reporting the results for linkage group (LG) 14 that harbors a major non-recombining area (supergene).

*Characterization of the supergene*

To investigate the potential supergene identified in the current study, we used SNP markers from queens (whole genome resequencing) and males (RADseq). We used BCFtools' (v.1.9) *isec* and *merge* functions to extract shared SNPs between the two datasets.

182 out of the resulting 1,064 SNP markers were used to visualize the genotypes of the males and queens at LG14 using VIVA (v.0.4.0) [45]. To further characterize the two haplotypes of the supergene, we performed a PCA on 69 SNP markers identified in the non-recombining area of LG14 (i.e. scaffolds 22, 23, 38, 42, 48, and parts of 6 and 14) in datasets of males and queens (Figure 3C and Supplemental file, Figure S9).

To investigate recombination suppression in the supergene region, we used SNP data from six queens homozygous for the Sh allele (H104W, H33G, H34G, H38W, H73W and H98W) and six others homozygous for the Sp allele (P22P, P102R, P114W, P109R, P34R, P84P), assigned based on visual inspection of their genotypes (Supplemental file, Figure S7). We first performed a VCF lift over (on scaffolds 14, 22, 23, 38, 42, 48 and 6 into LG14) using flo [46] to generate a chain file and picard's *LiftoverVcf* tool (v.2.20.0) to perform the lift over. We then used VCFtools (v.0.1.16) [14] to only keep bi-allelic variants with a MAF ≥ 0.2 and to thin the sites to help plotting and visualization. We next used LDheatmap (v.1.0-4) [47] to generate heatmap plots to visualize pairwise linkage disequilibrium (LD). The lack of recombination between Sh and Sp was verified by calculating pairwise linkage disequilibrium in a pool of twelve queens (six homozygous for Sh and six homozygous for Sp). Further, we used another six heterozygous (Sh/Sp) queens (H118W, H123W, H144W, H46G, H89W and P99P) to confirm the lack recombination between Sh and Sp (Supplemental file, Figure S10).

To investigate the genomic architecture of the supergene, we used BEDOPS' *bedmap* program [48] to estimate TE as well as gene content across LG14 in 100 kb sliding windows based on annotations produced as described above.

To explore whether any of the transcripts previously shown to be associated with aggression of queens depending on their social context [49] belong to the supergene, we aligned the assembled transcripts to the Pcal2 reference genome using GMAP [50]. We only retained the top hits with at least 70% query coverage, yielding 7303 transcripts (out of 7890) unambiguously aligned to the genome. We then used BEDtools' *intersect* function (v.2.29.2) [34] to extract transcripts that mapped to the supergene region (Dataset S2).

*<u>Dating the supergene formation</u>*

We approximately estimated the divergence between Sh and Sp by computing the rCCR, using MSMC2 [22] on four single individual queens with the Sh (H104W, H38W, H73W and H98W) and the Sp alleles (P114W, P50R, P84P and P97R). We limited the analysis to the supergene region (scaffolds 22, 23 and part of scaffold 6; larger than 1 Mb [51]) and ran 500 bootstraps by randomly sampling 2 Mb each time. Further, we performed a phylogenetic analysis of Sh and Sp alleles with

*P. subnitidus* as an outgroup species using RAxML [52]. Sequencing data of one *P. subnitidus* individual was generated and processed as described above. Joint SNP calling (with the other 35 queens sequenced in this study) was performed using GATK [12]. Next, 33,424 SNPs (only bi-allelic, polymorphic and missingness < 0.2) at the supergene region (scaffolds 22, 23, 38, 42, 48 and parts of scaffolds 6 and 14) were extracted and an alignment file was generated based on these SNPs for six queens homozygous for the Sh allele (H104W, H33G, H34G, H38W, H73W and H98W), six others homozygous for the Sp allele (P22P, P102R, P114W, P109R, P34R, P84P) and one *P. subnitidus* sample. The alignment file was then used to construct a maximum-likelihood (ML) tree using RAxML [52] under the GTRGAMMA model with bootstrapping.

We then used a fossil-calibrated dated phylogeny [53] to date the split between the Sh and Sp haplotype groups using a phylogenetic approach. The Ward et al. [53] phylogeny dated the split between *P. vermiculatus* and *P. imberbiculus* at 18 (95% highest probability density (HPD) ±8) MYA. This is an upper bound on the speciation date of *P. californicus* and *P. subnitidus* according to the most recent *Pogonomyrmex* phylogeny [54]. By comparing the *P. subnitidus* branch length (1.273) and the average path length from the split between Sh–Sb to the Sh leaves (0.0325) (haplometrotic queen sequences), we calculated a ratio of 1:39 between the speciation time of *P. subnitidus* and *P. californicus*, and the Sh–Sb haplotypes divergence time. Applying this ratio to the dating reference (i.e. 18±8 MYA), we obtained 0.46 (95% HPD ±0.20) MYA as an upper bound for the divergence between the two haplotype groups, which is consistent with the MSMC2 analysis. We note that the confidence in the age inference is strengthened by the agreement between two distinct approaches: the inference by MSMC2 is based on a coalescent model, which is a distinct approach from the phylogenetic dating approach. The former approach depends on estimates of mutation rate and generation time, whereas the latter depends on fossil calibration points. The tree figure was produced using interactive Tree Of Life (iTOL) [55].

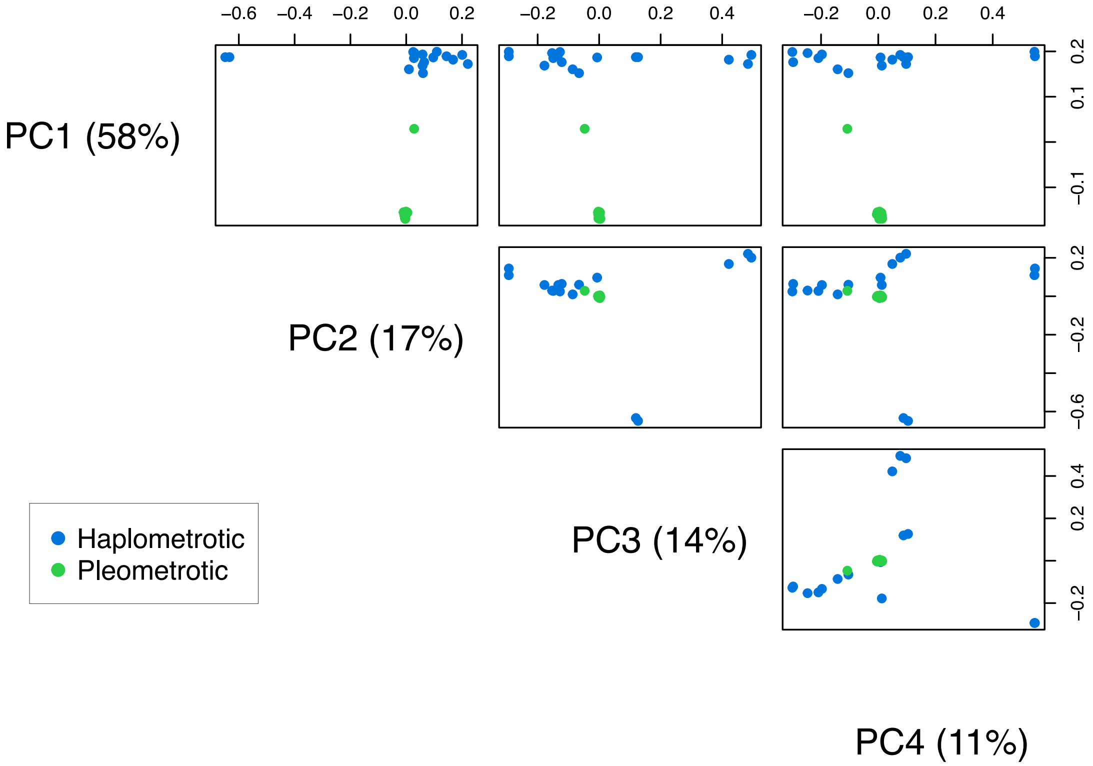


**Figure S1:** Principal Component Analysis based on 314,756 genome-wide SNPs of queens of the pleometrotic and haplometrotic populations. The scatterplot matrix shows the four eigenvectors (i.e. principal components). Note that only the haplometrotic queens vary along PCs 2, 3 and 4.

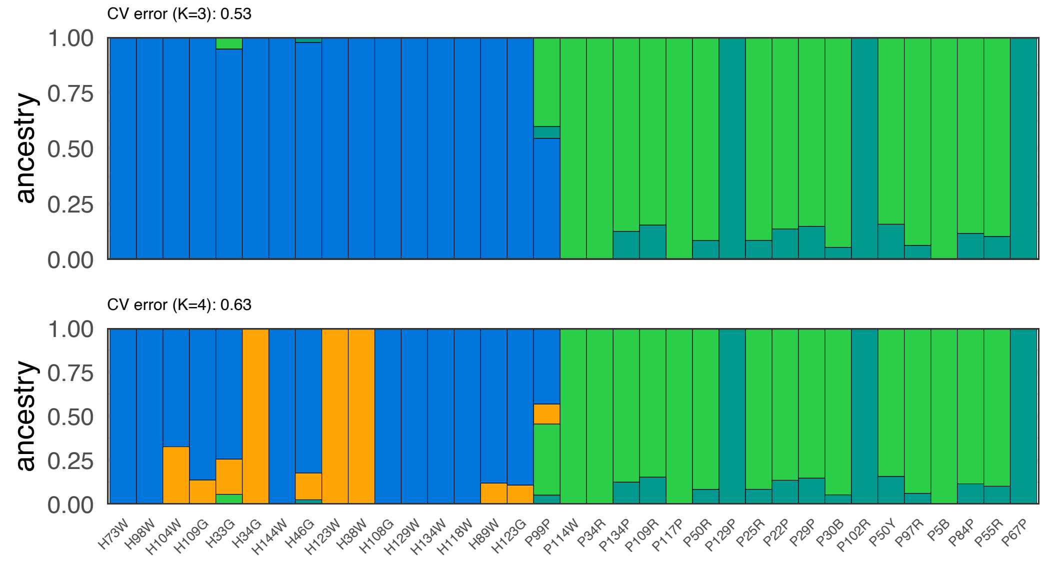


**Figure S2:** Admixture patterns among *P. californicus* populations using ADMIXTURE at K = 3 and K = 4. For each value of k, the cross-validation error rate is presented.

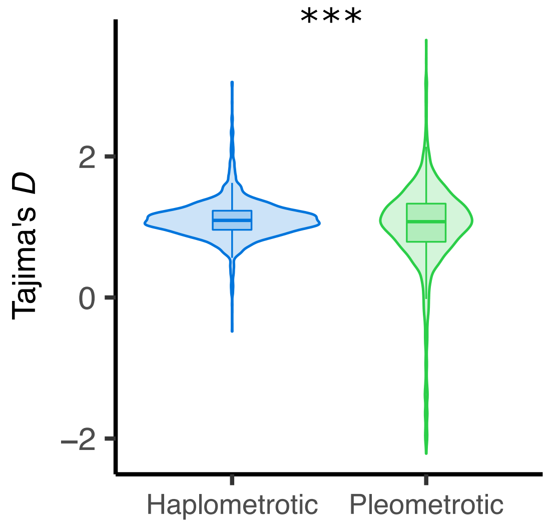


**Figure S3.** Tajima's *D* estimates calculated in 100 kb non-overlapping windows across the genome and averaged for the pleometrotic (green) and haplometrotic (blue) populations. ***: $P < 0.0001$.

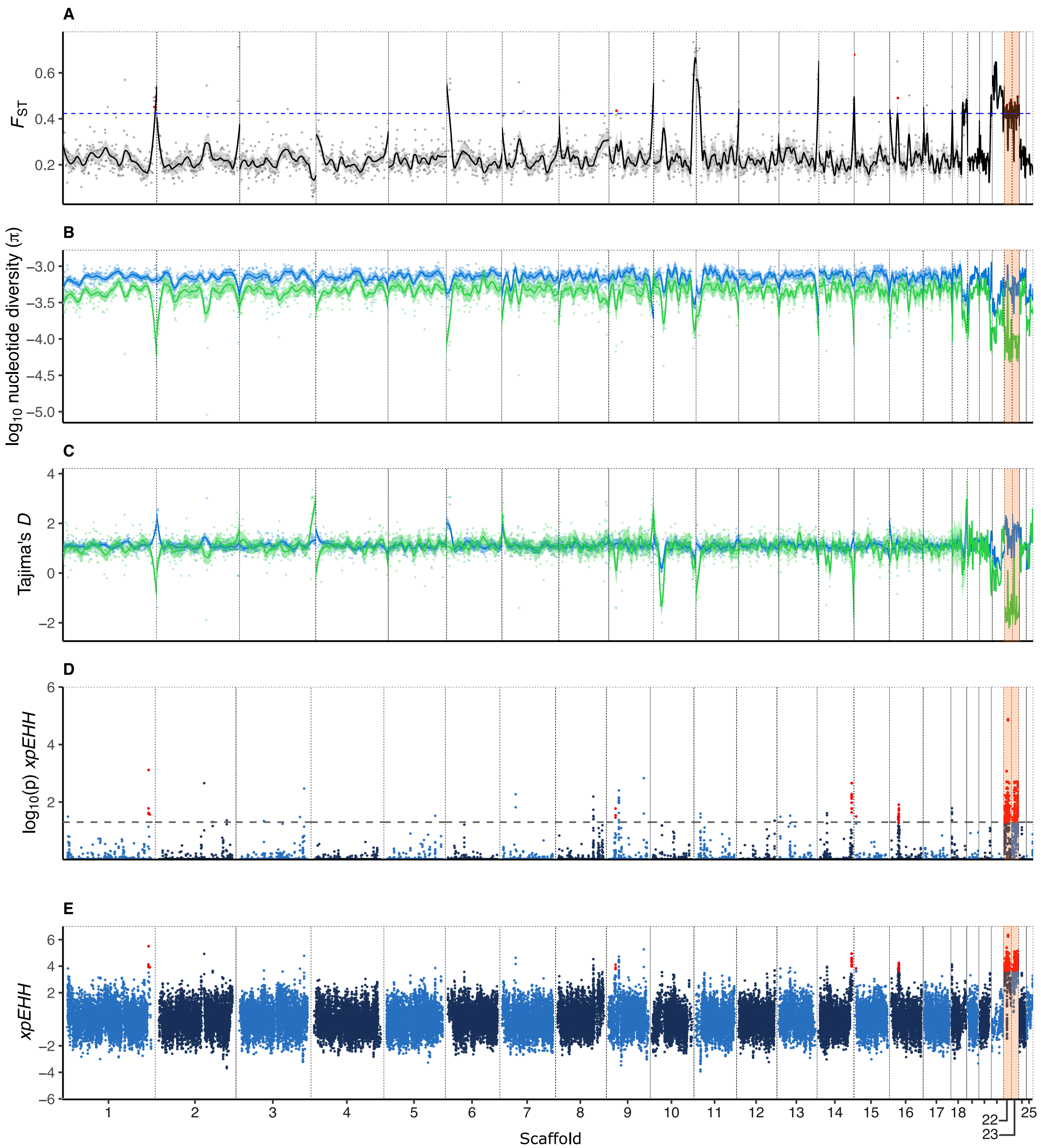


**Figure S4.** Genome-wide genetic differentiation ($F_{ST}$), nucleotide diversity (π), Tajima's *D* and *xpEHH* scores in the 25 largest scaffolds of the *P. californicus* genome. (A) Genetic differentiation ($F_{ST}$) between haplometrotic and pleometrotic queens. Blue dashed line indicates the 5% cutoff. (B) Nucleotide diversity ($\pi$) and (C) Tajima's *D* in the haplometrotic (blue) and pleometrotic (green) populations. All estimates were calculated in 100 kb non-overlapping windows. Lines and shaded areas are means and 95% confidence intervals, respectively. (D) log-scaled P-values and (E) *xpEHH* scores associated with SNPs across the 25 scaffolds. Black dashed line represents threshold of significance for outlier SNPs ($p < 0.05$). Red dots highlight windows ($F_{ST}$) and SNPs (*xpEHH*) showing signatures of selection in both analyses. Scaffolds 22 and 23 containing the putative supergene are highlighted in orange.

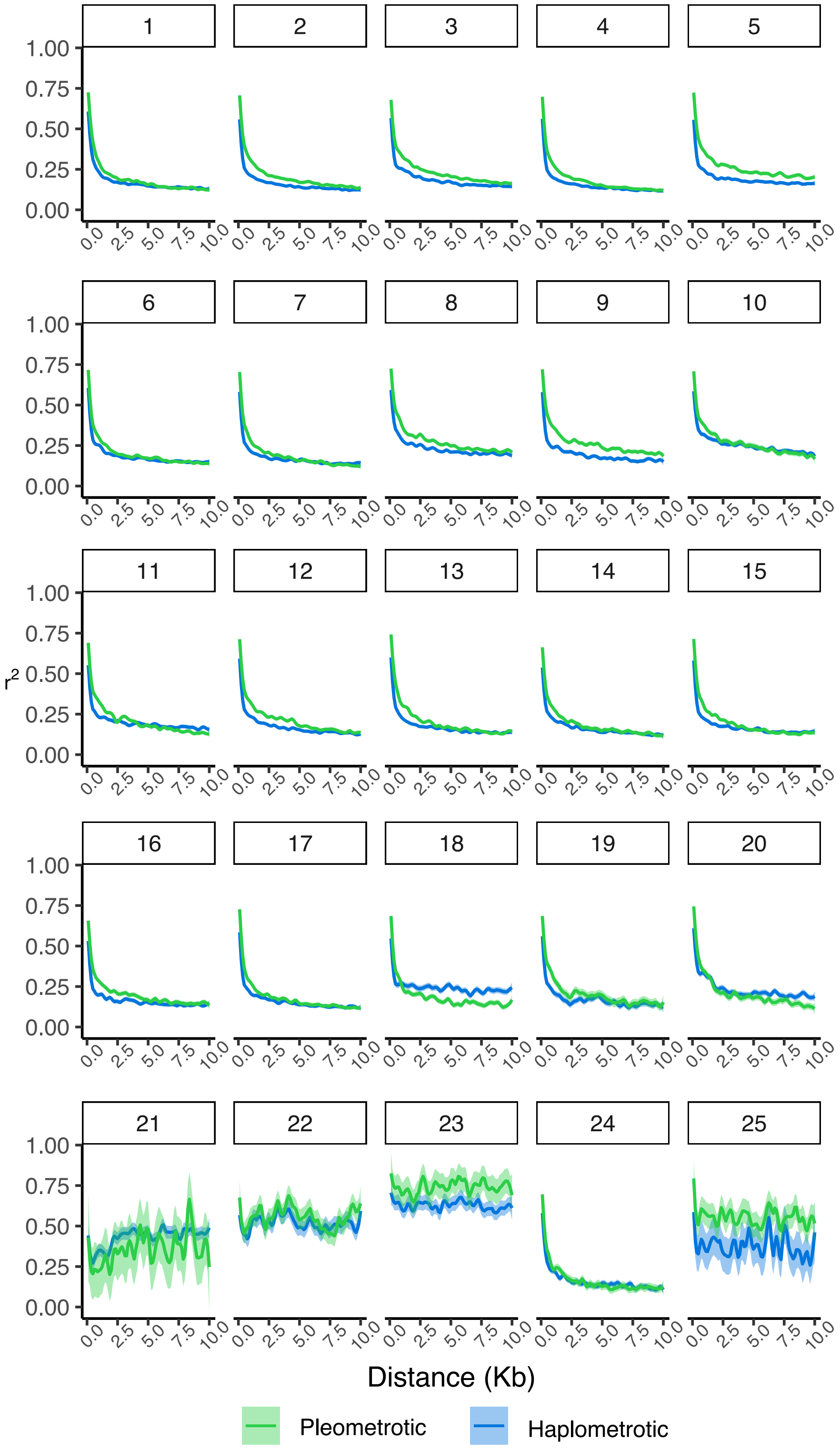


**Figure S5.** Linkage disequilibrium (LD) decay displayed for each of the 25 largest scaffolds in the two *P. californicus* populations. LD as measured by pairwise $r^2$ is represented on the y-axis and distance between pairs of markers within 10 kb on the x-axis. Lines and shaded areas are means and 95% confidence intervals, respectively.

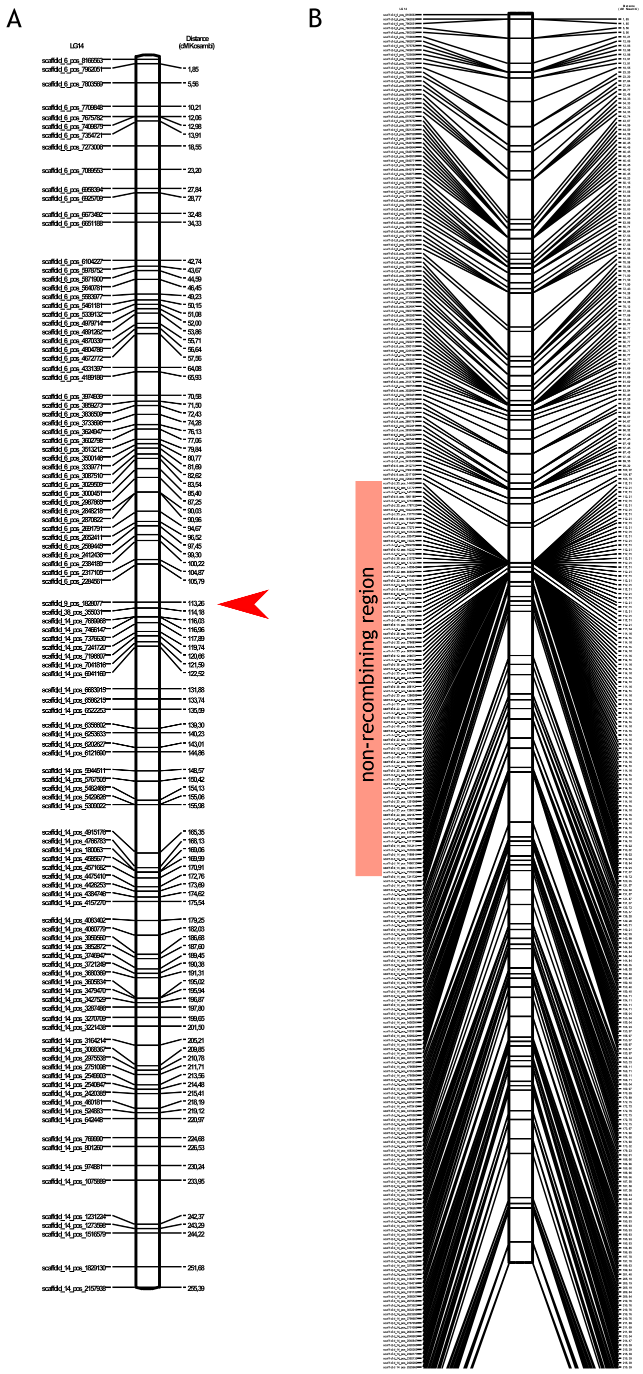


**Figure S6.** Linkage group (LG) 14 (253.38 cM). Marker names are given on the left and genetic distance in cM (Kosambi) are given on the right sides. Both figures represent the same LG group, panel B includes all 339 markers and shows the non-recombining area. Panel A shows a “skeleton linkage group” where overlapping markers at the same position are grouped and represented only by a single marker name (i.e. the non-recombining region is represented by a single marker; thick red arrowhead).

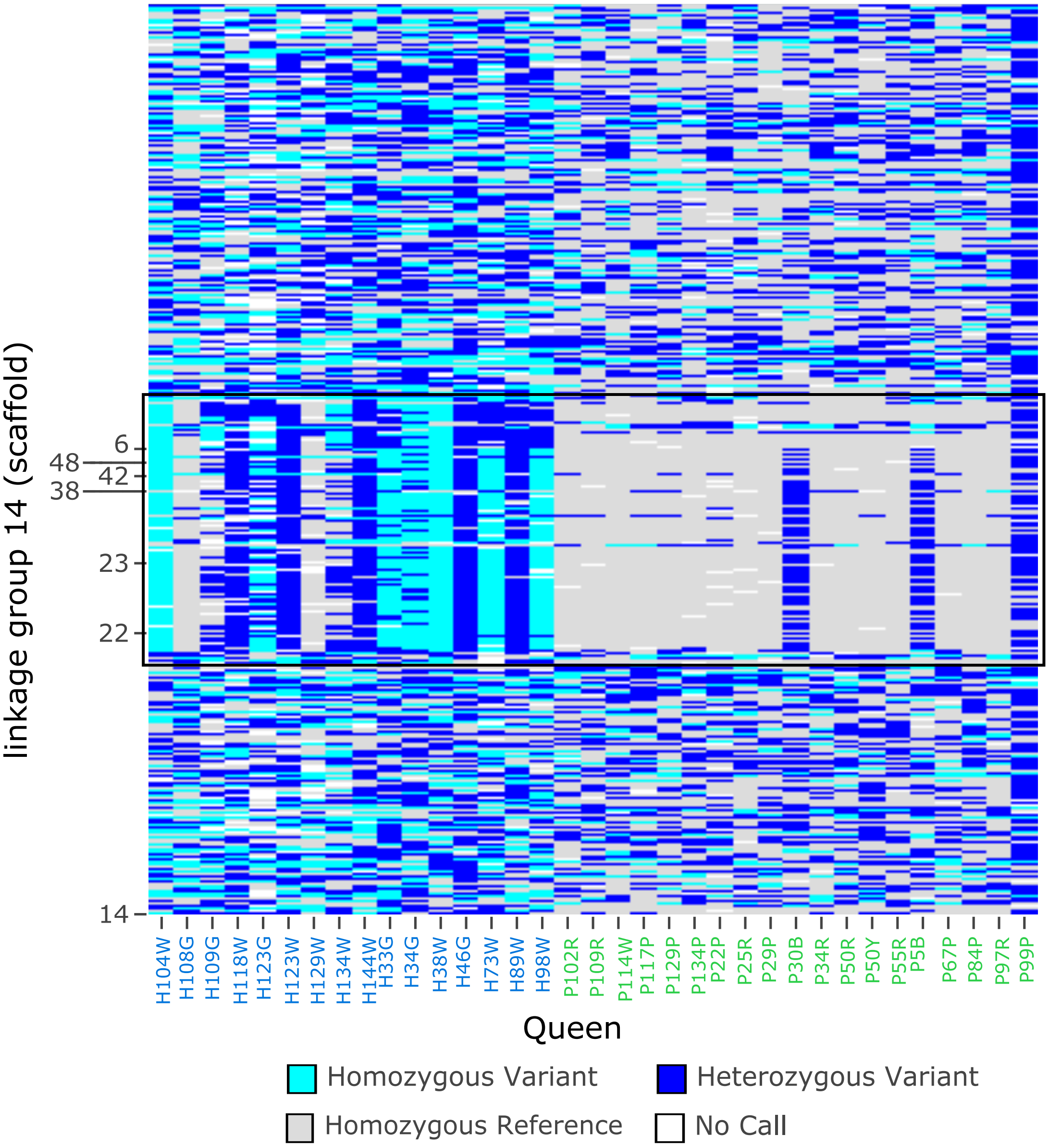


**Figure S7.** Genomic architecture at linkage group (LG) 14 harboring the supergene in *P. californicus*. The heatmap displays the genotypes of the 35 queen samples from the pleometrotic and haplometrotic populations. Rows represent a total of 5,363 SNPs (thinned out of 68,643 SNPs for ease of visualization) with a MAF ≥ 0.2, spanning seven scaffolds of LG14. Columns represent individual queens grouped by population (pleometrotic and haplometrotic). Black rectangle delimits the non-recombining supergene region, spanning scaffolds 22, 23, 38, 42, 48 and parts of scaffolds 6 and 14. Queen P99P is an F1 hybrid collected from a pleometrotic population.

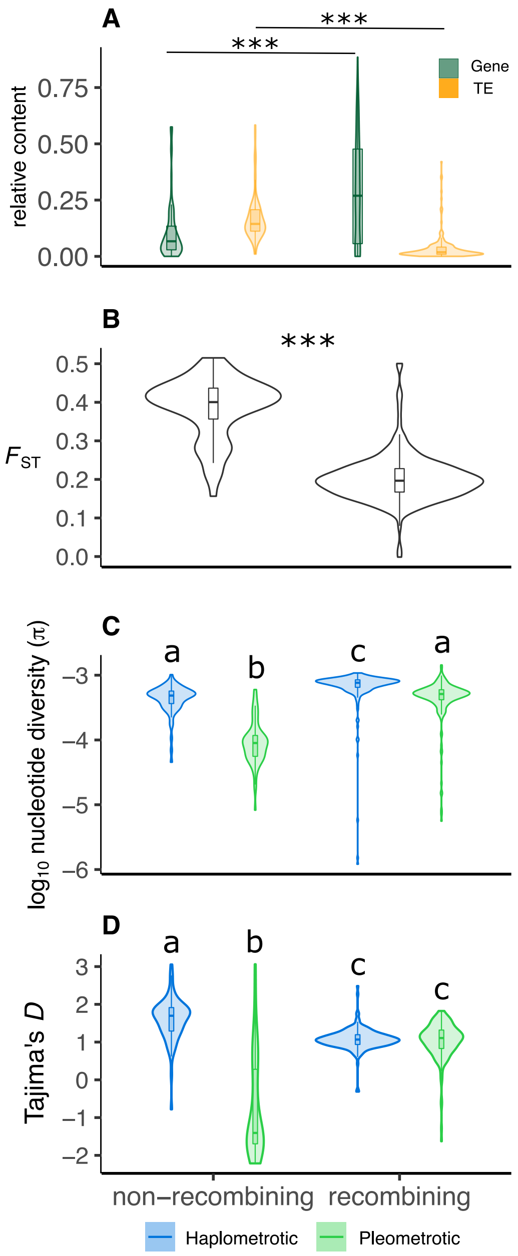


**Figure S8.** Genetic architecture of the supergene in *P. californicus*. (A) Violin plots showing exon and TE content within non-recombining regions (i.e. the supergene) and recombining regions of LG14. We performed a Kruskal-Wallis rank sum test ($\chi^2$= 342.74, df = 3, p < 2.2e-16), followed by pairwise Wilcoxon rank sum *post hoc* tests. (B) Genetic differentiation ($F_{ST}$), (C) nucleotide diversity ($\pi$) and (D) Tajima's *D* within recombining and non-recombining regions of LG14 in the pleometrotic and haplometrotic populations. For $F_{ST}$, we performed a Wilcoxon rank sum test (W = 12261, p < 2.2e-16). We performed a Kruskal-Wallis rank sum test for $\pi$ ($\chi^2$ = 254.77, df = 3, p < 2.2e-16) and Tajima's D ($\chi^2$ = 177.81, df = 3, p < 2.2e-16), followed by pairwise Wilcoxon rank sum *post hoc* tests. Different letters represent significant differences (p < 0.0001).

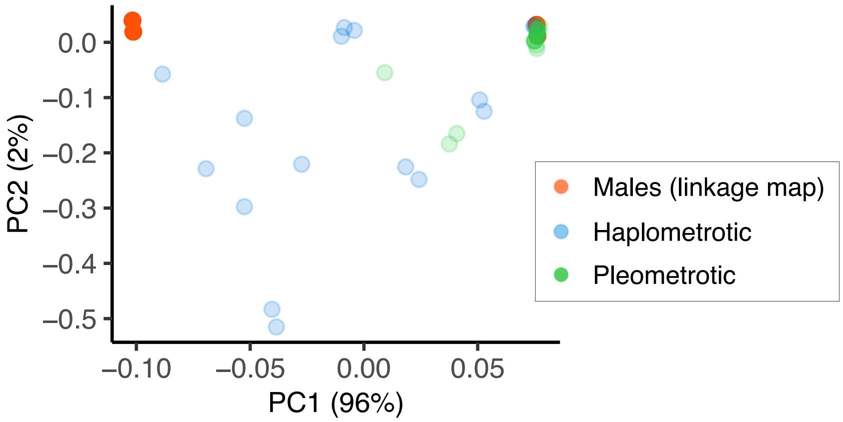


**Figure S9.** Principal Component Analysis based on 69 markers spanning the supergene region showing that the 108 male haplotypes are either Sh (negative PC1 scores) or Sp (positive PC1 scores). Diploid queens are either homozygous for the Sp allele (tightly clustered green points; most of pleometrotic queens) or heterozygous Sh/Sp (widely distributed blue points; most of the haplometrotic queens).

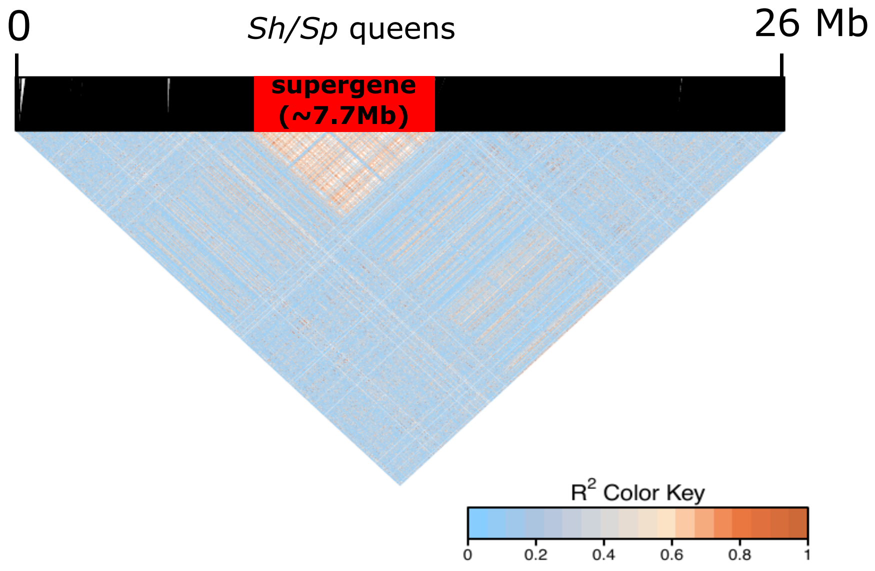


**Figure S10.** Linkage disequilibrium estimates ($r^2$) across linkage group (LG) 14 containing the supergene for six heterozygous queens (Sh/Sp). The red rectangle shows the region of no recombination between Sh and Sp. SNPs are ordered according to physical position on the chromosome after lift over. Black lines on top of the heatmap indicate SNP positions on the physical map.

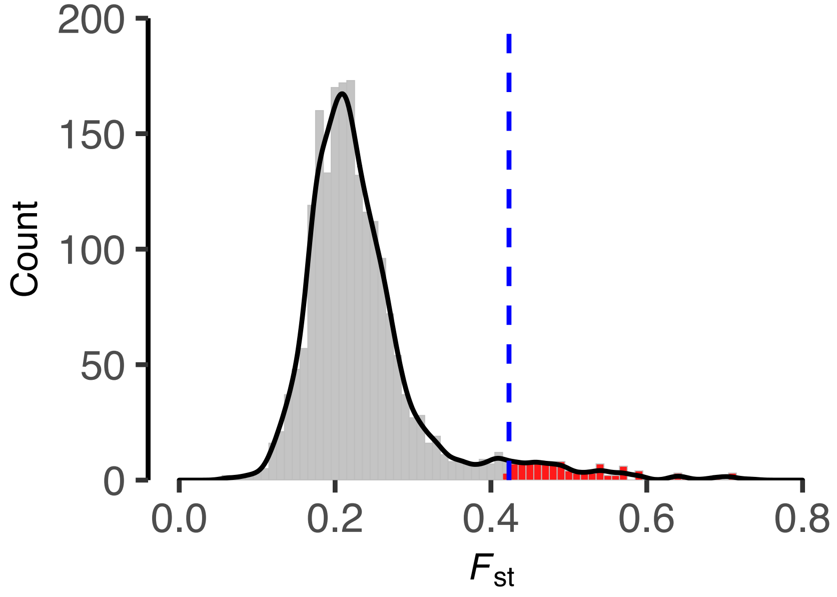


**Figure S11.** Distribution of pairwise genetic differentiation ($F_{ST}$) between haplometrotic and pleometrotic queens, estimated in 100 kb non-overlapping windows across the genome. Regions falling into the upper 5% tail of the distribution ($F_{ST}$ > 0.423, delimited by the blue dashed line) were considered candidates for divergent selection between populations.

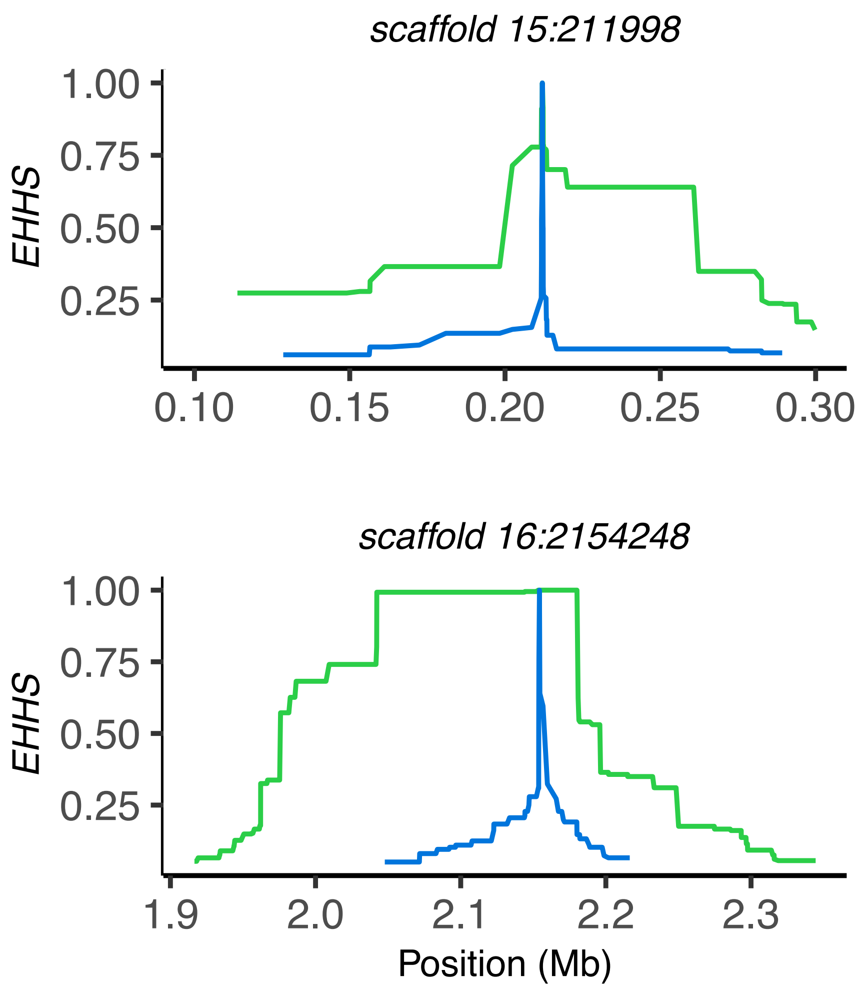


**Figure S12.** Signatures of site-specific extended haplotype homozygosity (*EHHS*) at two outlier SNPs in the pleometrotic (green), but not the haplometrotic population (blue) indicating recent selective sweeps. The position of each SNP is indicated at the top of each plot. Our six best candidate genes are found in these regions of elevated extended haplotype homozygosity scores.

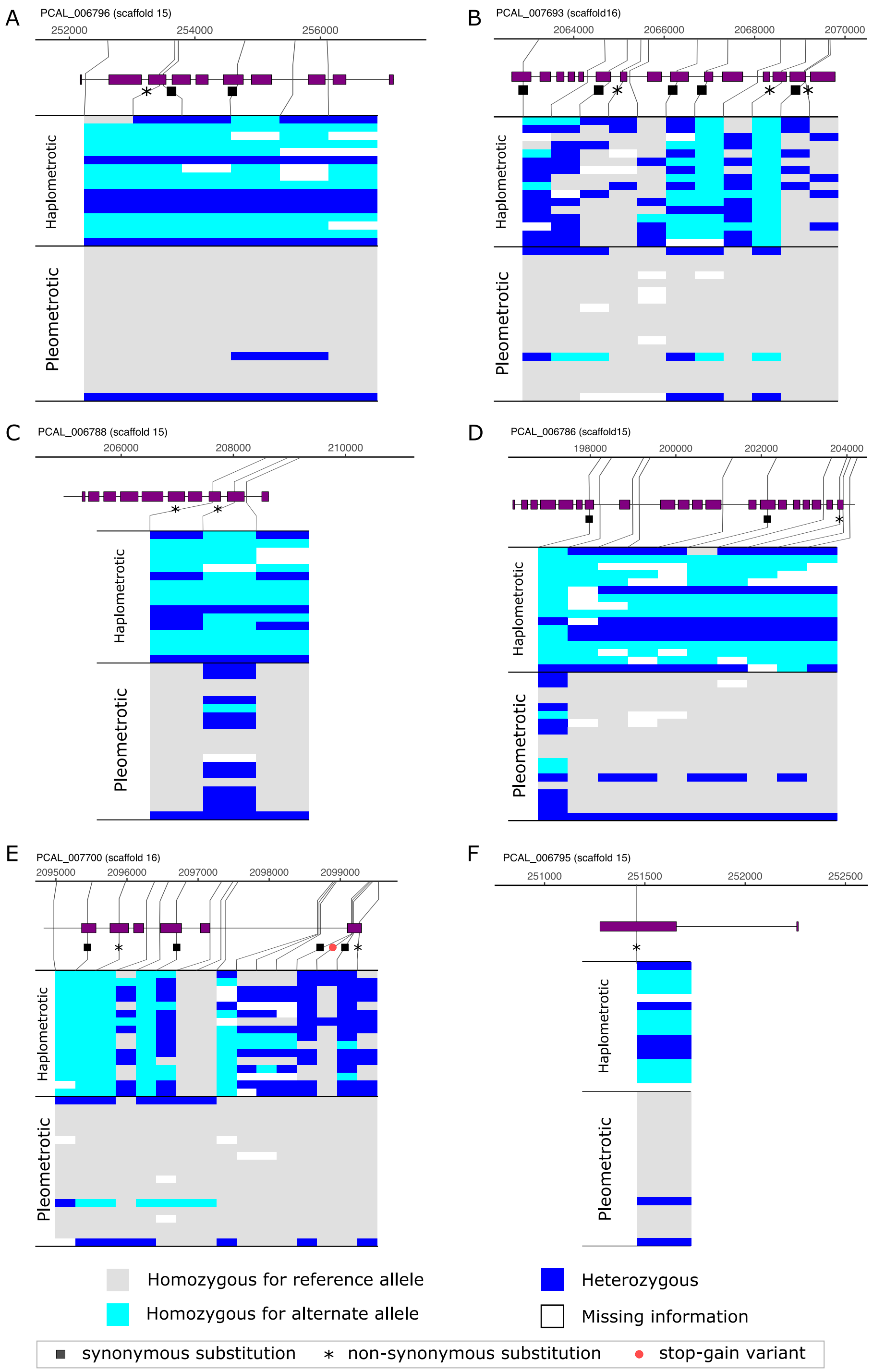


**Figure S13.** Genotypes of haplo- and pleometrotic queens for candidate genes under selection. Each plot shows SNP positions (top, solid lines), the affected features of the candidate gene (gene models in purple), and the genotypes for each queen in the two populations. Pleometrotic queens are almost all homozygous and fixed for the reference allele, unlike the haplometrotic queens that are more variable. Plots were produced using GenotypePlot [https://github.com/JimWhiting91/genotype_plot]. Note that the bottom row in the pleometrotic population displays the genotype of the hybrid P99P queen.

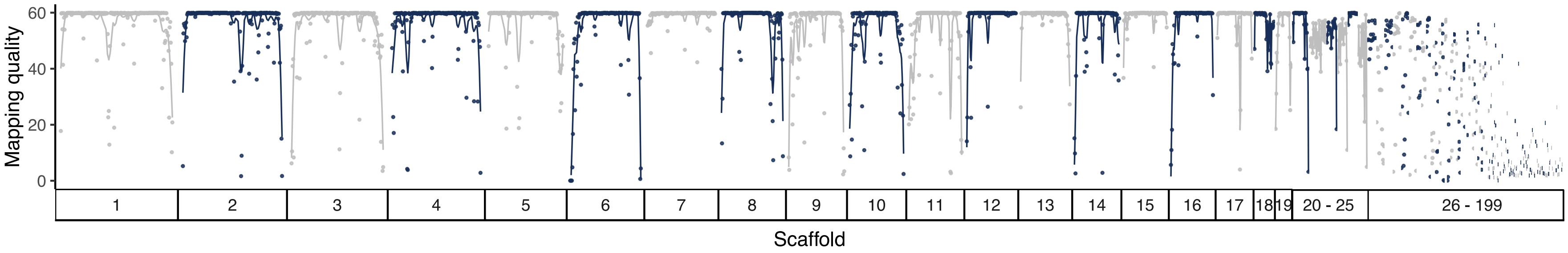


**Figure S14.** Mapping quality across the 199 scaffolds of the Pcal2 assembly. Dots represent average mapping quality in 100 kb window, calculated and averaged across the 35 queens sequenced in this study. Scaffolds 26-199 were excluded from all analyses due to fragmentation and low mapping quality.

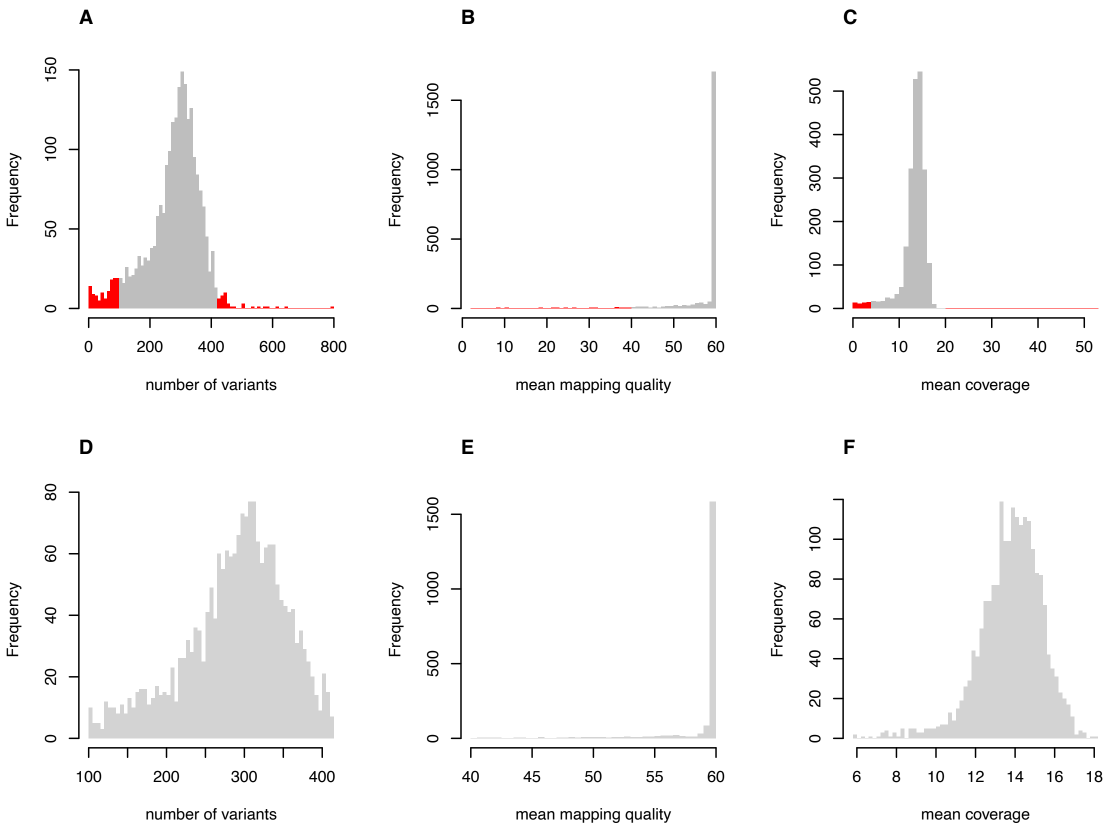


**Figure S15.** Quality control of non-overlapping genomic windows (100 kb) used to compute population genomic metrics. Number of variants, mapping quality and mean coverage per 100 kb window before (A, B, and C, respectively) and after (D, E, and F, respectively) applying cutoffs (see Material and Methods). Red bars indicate windows excluded from population genomic analyses.

**Table S1.** Quality control metrics and phenotypes of queens sequenced in this study. Sandwich assays were performed to phenotype the queens as singleton or in pairs (see Material and Methods). H: a single haplometrotic queen, P: a single pleometrotic queen, HH: the two queens are haplometrotic, PP: the two queens are pleometrotic and HP: a haplometrotic and a pleometrotic queen. The first letter in column nest type represents the sequenced queen, i.e. HP means that the sequenced queen was from the haplometrotic population and PH the inverse.

| Queen | No. reads | Percentage mapped | Mean coverage | Mapping quality | Insert size | Population | Year of sampling |
|---|---|---|---|---|---|---|---|
| H104W | 27215573 | 99.9 | 8 | 41 | 270 | haplometrotic | 2018 |
| H108G | 35400779 | 99.8 | 10 | 41 | 254 | haplometrotic | 2018 |
| H109G | 65438542 | 99.9 | 19 | 40 | 354 | haplometrotic | 2018 |
| H118W | 72316975 | 99.9 | 21 | 40 | 295 | haplometrotic | 2018 |
| H123G | 76030603 | 99.8 | 22 | 40 | 277 | haplometrotic | 2018 |
| H123W | 62340922 | 99.2 | 18 | 40 | 232 | haplometrotic | 2018 |
| H129W | 71925022 | 99.8 | 21 | 40 | 286 | haplometrotic | 2018 |
| H134W | 59749398 | 99.9 | 17 | 40 | 267 | haplometrotic | 2018 |
| H144W | 33707630 | 99.9 | 10 | 41 | 274 | haplometrotic | 2018 |
| H33G | 41096388 | 99.9 | 12 | 40 | 217 | haplometrotic | 2018 |
| H34G | 35144716 | 98.9 | 10 | 40 | 247 | haplometrotic | 2018 |
| H38W | 41023193 | 99.9 | 12 | 41 | 240 | haplometrotic | 2018 |
| H46G | 45041325 | 99.8 | 13 | 40 | 203 | haplometrotic | 2018 |
| H73W | 66932264 | 99.9 | 19 | 40 | 343 | haplometrotic | 2018 |
| H89W | 44369950 | 99.2 | 13 | 41 | 215 | haplometrotic | 2018 |
| H98W | 55388781 | 99.8 | 17 | 40 | 278 | haplometrotic | 2018 |
| P99P | 62941605 | 99.8 | 18 | 40 | 250 | pleometrotic | 2018 |
| P102R | 44321548 | 99.9 | 13 | 41 | 274 | pleometrotic | 2018 |
| P109R | 54702700 | 99.7 | 16 | 40 | 256 | pleometrotic | 2018 |
| P114W | 46302465 | 99.9 | 14 | 41 | 210 | pleometrotic | 2018 |
| P117P | 56580092 | 99.9 | 16 | 40 | 250 | pleometrotic | 2018 |
| P129P | 37104772 | 99.9 | 11 | 41 | 219 | pleometrotic | 2018 |
| P134P | 60686603 | 99.8 | 17 | 40 | 253 | pleometrotic | 2018 |
| P22P | 58042696 | 99.7 | 17 | 40 | 234 | pleometrotic | 2018 |
| P25R | 46013864 | 99.9 | 14 | 40 | 247 | pleometrotic | 2019 |
| P29P | 32986286 | 99.9 | 10 | 41 | 185 | pleometrotic | 2018 |
| P30B | 43225857 | 99.9 | 13 | 40 | 221 | pleometrotic | 2019 |
| P34R | 40153297 | 99.9 | 12 | 40 | 198 | pleometrotic | 2018 |
| P50R | 50428119 | 99.8 | 15 | 40 | 240 | pleometrotic | 2019 |
| P50Y | 37180800 | 99.9 | 11 | 41 | 191 | pleometrotic | 2018 |
| P55R | 43176568 | 97.8 | 13 | 41 | 212 | pleometrotic | 2018 |
| P5B | 31292850 | 99.9 | 9 | 40 | 225 | pleometrotic | 2018 |
| P67P | 35099092 | 99.9 | 10 | 41 | 220 | pleometrotic | 2018 |

| | | | | | | | |
|---|---|---|---|---|---|---|---|
| P84P | 41853659 | 99.9 | 12 | 40 | 228 | pleometrotic | 2018 |
| P97R | 41252327 | 99.9 | 12 | 40 | 245 | pleometrotic | 2018 |

**Table S2.** Genomic coordinates and sizes of regions belonging to the supergene on LG14 including whole scaffolds 22, 23, 38, 42, 48, and parts of scaffolds 6 and 14.

| Scaffold | Start | End | Size (Mb) |
|---|---|---|---|
| scaffold_14 | 7736831 | 8322736 | 0.59 |
| scaffold_22 | 0 | 1862986 | 1.86 |
| scaffold_23 | 0 | 1832116 | 1.83 |
| scaffold_38 | 0 | 564213 | 0.56 |
| scaffold_42 | 0 | 471449 | 0.47 |
| scaffold_48 | 0 | 387552 | 0.39 |
| scaffold_6 | 0 | 2020227 | 2.02 |

**Table S3.** Results for Hardy–Weinberg calculations at the supergene locus for the pleometrotic and haplometrotic populations. IDs of queen assigned to each genotype (Sh/Sh, Sp/Sp or Sh/Sp) are indicated for each population. Genotypes were assigned based on visual inspection of the supergene region (see Material and Methods).

| **Pleometrotic population** | | | | |
|---|---|---|---|---|
| | observed | expected | $\chi^2$ | Queen ID |
| Sp/Sp | 16 | 16.1 | 0.00087 | P102R, P109R, P114W, P117P, P129P, P134P, P22P, P25R, P29P, P34R, P50R, P50Y, P55R, P67P, P84P, P97R |
| Sh/Sp | 3 | 2.76 | 0.0203 | P30B, P5B, P99P |
| Sh/Sh | 0 | 0.118 | 0.118 | |
| sum | 19 | 19.0 | 0.1396 | |
| Allele frequency (Sp) | 92.1% | | | |
| Allele frequency (Sh) | 7.9% | | | |
| **$\chi^2$ for 1 degree of freedom at α = 0.05** | | | | 3.841 |
| | | | | 0.1396 < 3.841 |
| **P-value** | | | | 0.7087 |

| **Pleometrotic population** | | | | |
|---|---|---|---|---|
| | observed | expected | $\chi^2$ | Queen ID |
| Sp/Sp | 2 | 2.25 | 0.028 | H108G, H129W |
| Sh/Sp | 8 | 7.50 | 0.03 | H109G, H118W, H123G, H123W, H134W, H144W, H46G, H89W |
| Sh/Sh | 6 | 6.25 | 0.010 | H104W, H33G, H34G, H38W, H73W, H98W |
| sum | 16 | 16.0 | 0.0711 | |
| Allele frequency (Sp) | 37.5% | | | |
| Allele frequency (Sh) | 62.5% | | | |
| **$\chi^2$ for 1 degree of freedom at α = 0.05** | | | | 3.841 |
| | | | | 0.0711 < 3.841 |
| **P-value** | | | | 0.7897 |

**Table S4.** Genomic coordinates of regions showing patterns of positive selective sweeps in the pleometrotic population, identified by our two selection scan approaches ($F_{ST}$ and *xpEHH*).

| scaffold | start | end |
|---|---|---|
| scaffold_1 | 19964834 | 20100000 |
| scaffold_1 | 20277427 | 20477427 |
| scaffold_9 | 1902145 | 2000000 |
| scaffold_14 | 8051123 | 8200000 |
| scaffold_15 | 200000 | 300000 |
| scaffold_16 | 2054248 | 2200000 |
| scaffold_22 | 100000 | 200000 |
| scaffold_22 | 951559 | 1000000 |
| scaffold_22 | 1100000 | 1151559 |
| scaffold_22 | 1400000 | 1500000 |
| scaffold_23 | 200000 | 386785 |
| scaffold_23 | 633946 | 800000 |
| scaffold_23 | 1107655 | 1200000 |
| scaffold_23 | 1300000 | 1310010 |
| scaffold_23 | 1417040 | 1500000 |
| scaffold_23 | 1600000 | 1617040 |
| scaffold_23 | 1647159 | 1700000 |

**Dataset S1 (separate file).** Detailed report on *Pogonomyrmex californicus* genome sequencing, assembly and annotation.

**Dataset S2 (separate file).** List of transcripts found in the supergene region on LG14 that were shown previously to be differentially expressed between queens of the two populations in the context of origin (pleometrotic/haplometrotic), behavior (aggressive/nonagressive) or social environment. See [49] for description of the social context.

**Dataset S3 (separate file).** Top 5% highly differentiated regions between the haplometrotic and pleometrotic populations. Genomic position, number of SNPs, $F_{ST}$, π, Tajima's *D*, mean coverage, and mean mapping quality (MQ) are displayed for each candidate region.

**Dataset S4 (separate file).** Outlier SNPs showing clear peaks of *xpEHH* ($p<0.05$) using the Benjamani & Hochberg method for FDR correction. Positive values in the *xpEHH* column indicate selection in the pleometrotic population, while negative values indicate selection in the haplometrotic population. SNPs within 100 kb of each other were grouped in the same cluster (32 clusters indicated by the cluster column).

**Dataset S5 (separate file).** Candidate genes showing signatures of selection in the pleometrotic population identified by our two selection scan approaches ($F_{ST}$ and *xpEHH*). 34 likely functional genes were kept as they are not encoding TE proteins, are expressed, or homologous to other known insect proteins (see Material and Methods).

**Dataset S6 (separate file).** Phylogenetic tree used to date the supergene, produced by RAxML using the GTRGAMMA model, in Newick format.
