## Supplementary material for "Evolutionary genomics of socially polymorphic populations of *Pogonomyrmex californicus*": Dataset S1

### *Pogonomyrmex californicus* genome sequencing, assembly and annotation.

#### DNA extraction and library prep

We generated a single flowcell worth of minION data for the de novo genome assembly of the pleometrotic population of *Pogonomyrmex californicus*.

We sampled three pupae from colony **Pcal18-02** in liquid nitrogen for high-molecular weight DNA extraction using an phase-separation protocol, optimized for DNA extraction from ants (see *Materials and Methods*). We used 2 µg of HMW DNA for library preparation with the Ligation Sequencing Kit (Oxford Nanopore) **SQKLSK109**.

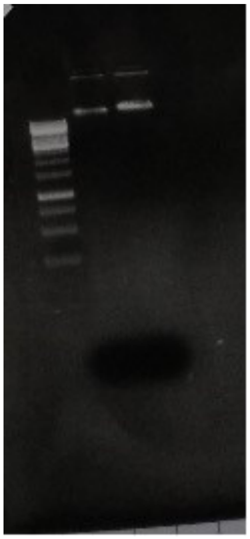

***Figure 1****: Agarose gel analysis of HMW DNA (lane 1) and the minION library (lane 2)*

The library was loaded to a minION sequencer and sequenced for 48 h. After basecalling with guppy (v4.0.14) we retrieved 1,650,688 reads (mean read length 4.9 kb) with a mean read quality of 13.4. Read length N50 was 18.5 kb and a total of 8.107 GB of data was generated.

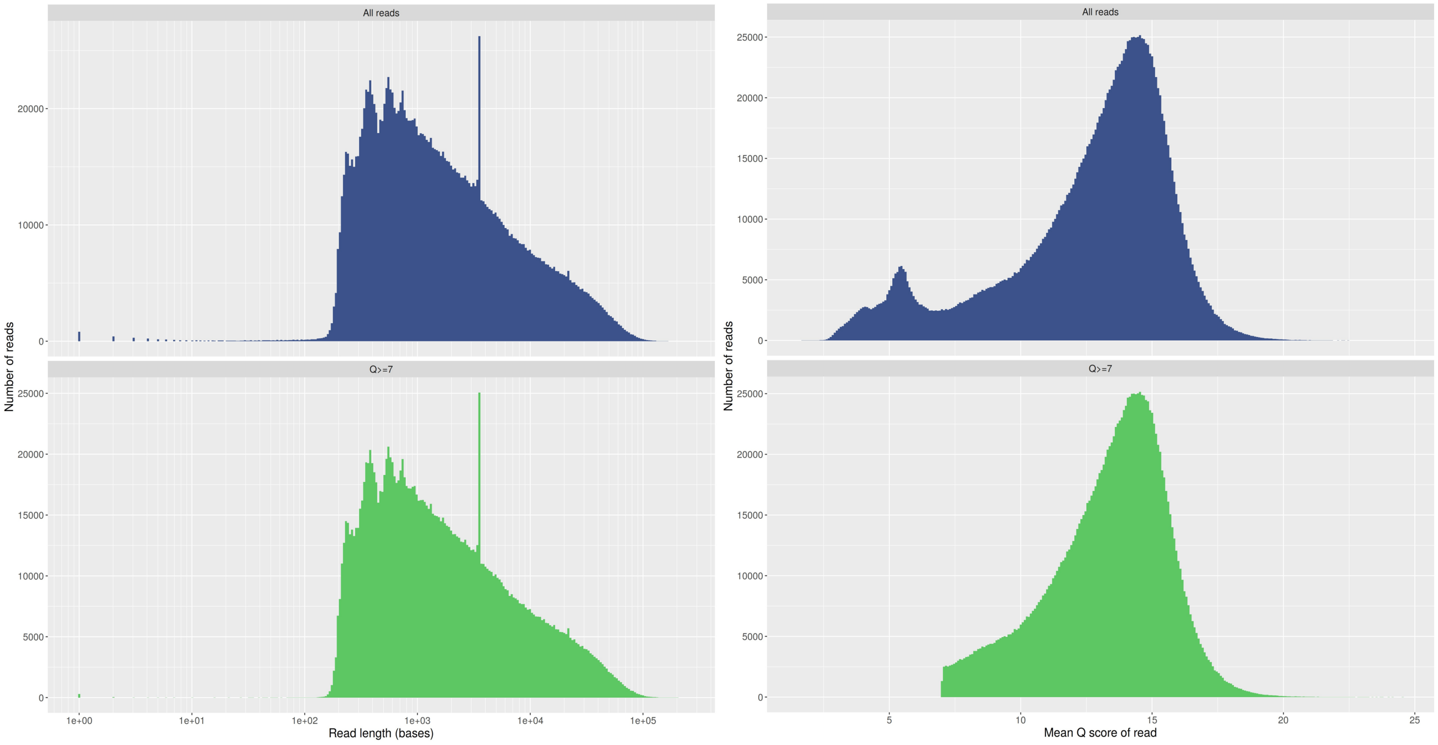


***Figure 2***: *Raw sequencing data output from the minION, the upper plots (in blue) show data for all reads and the lower plots (in green) show data for reads with a mean quality score above 7. Left plots: distribution of read length. Right plots: distribution of read mean.*

#### Raw data processing

We used PoreChop (v0.2.4) to remove adapter contaminations from minION reads before further filtering the data with FiltLong (v0.2.0). Filtlong allows selection of high quality reads based on comparison with short-read data of the same species. We sampled paired-end Illumina data of *Pogonomyrmex californicus* queens from for FiltLong from the following samples.

| Fastq | Sample |
|---|---|
| DNA_11_S2_R1_001.fastq.gz | DNA_11 |
| DNA_13_S4_R1_001.fastq.gz | DNA_13 |
| DNA_19_S10_R1_001.fastq.gz | DNA_19 |

We ran FiltLong using the following settings: filtlong –split 500 –min_length 3000 –keep_percent 90 –target_bases 7300000000 –trim –illumina_1 $illSample $nanoporeReads
Assuming a genome size of 240 Mb, we aimed to retrieve ~30 X long-read coverage for genome assembly.

#### Genome assembly

We assembled the genome from the ~30x Nanopore data using the following strategy. We created one assembly using `CANU (v2.0)` and a second assembly using `wtdbg2 (v2.5)` . Both assemblies were processed and optimized similarly, as follows: After contig assembly with CANU and wtdbg2, respectively, we used SSPACE for scaffolding with the long-read data. After gap filling with LR_closer, we used ntedit for a first round of polishing with long-reads. WE then used short-read, short-insert Illumina data for scaffolding with SSPACE before three rounds of polishing with Pilon.

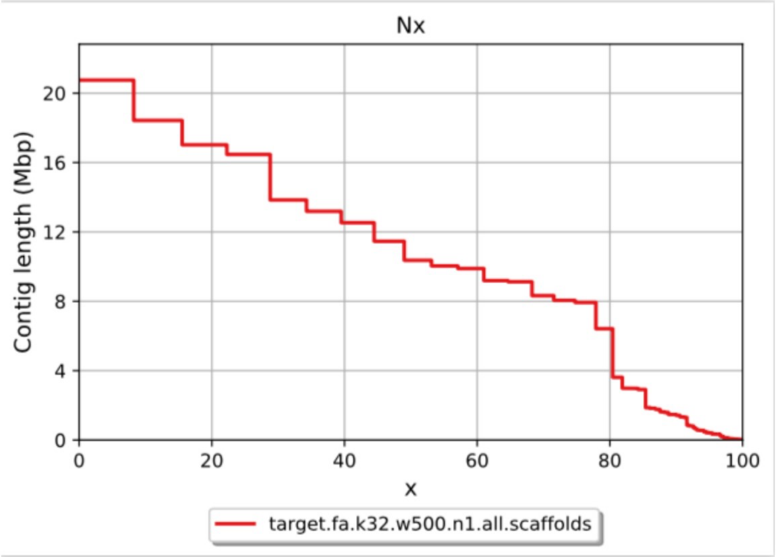


***Figure 3:*** *Nx plot for the final super-scaffolded and joint assembly for P. californicus*

The polished assembly was then super-scaffolded with publicly available 10X Chromium data for *P. californicus*
(SRA Accession SRX8051306) using `scaff10x` and subsequent correction with `break10x` . We used an inhouse, blastn-based pipeline to detect and remove bacterial contaminations from the assemblies.

Finally, we combined the CANU-based and the wtdbg-based assemblies with `ntjoin` , using the CANU-based assembly as reference. The joint assembly was filtered for redundancy using `funannotate clean` (v1.7.1), removing four scaffolds.

This produced a final assembly of the following quality:

| N50 | # scaffolds | largest scaffold | total size | # Ns | BUSCO |
|---|---|---|---|---|---|
| 10.36 MB | 199 | 20.75 MB | 252.297 MB | 337161 | C:98.5%[S:97.8%,D:0.7%],F:1.1%,M:0.4%,n:4415 |

Analysis with `BUSCOv3` (vs. hymenoptera_odb9) confirmed that this is a high-quality, nearly complete assembly (98.5% complete BUSCOs) with very low levels of duplication (0.7% duplicated BUSCOs).

#### Repeat annotation

We annotated repeats and transposable elements in the genome using a combination of de-novo, structural and homology-based approaches.

We ran `RepeatModeler2` and `EDTA` on thse genome assembly to generate de-novo repeat libraries. Libraries from both tools were combined and redundancies removed with `CD-HIT` . We classified repeats using a combination of `PasteClassifier` , `DeepTE` and `TEsorter` . We removed putative host proteins from the library by blasting all unclassified repeats against uniprot_arthropoda (uniprot-arthropoda+6656 downloaded 27.04.2018) with `blastx -soft_masking false -evalue 1e-10` .

We combined our de-novo library with public resources from RepBase (RepBase25.04.invrep.fa) and Dfam (Dfam3.0.arthropod.fa). Finally, using this combined library, we annotated the genome with RepeatMasker (v.open-4.0.7) with `-s -gff -a -inv -small -excln -cutoff 250 -nolow` . The produced annotation was subsequently improved further with `One-code-to-find-them-all` .

```
==================================================
file name: Pcal.genome.fa
sequences:           199
total length:  252297138 bp  (251960218 bp excl N/X-runs)
GC level:         37.13 %
bases masked:   57418338 bp ( 22.79 %)
==================================================
               number of      length   percentage
               elements*    occupied  of sequence
--------------------------------------------------
SINEs:              168        26006 bp    0.01 %
      ALUs            0            0 bp    0.00 %
      MIRs            0            0 bp    0.00 %

LINEs:              936       579803 bp    0.23 %
      LINE1           0            0 bp    0.00 %
      LINE2           3           45 bp    0.00 %
      L3/CR1          0            0 bp    0.00 %

LTR elements:     11520      5992778 bp    2.38 %
      ERVL            0            0 bp    0.00 %
      ERVL-MaLRs      0            0 bp    0.00 %
      ERV_classI    279        38288 bp    0.02 %
      ERV_classII     0            0 bp    0.00 %

DNA elements:     57365     11083347 bp    4.40 %
     hAT-Charlie     22        16393 bp    0.01 %
     TcMar-Tigger     0            0 bp    0.00 %

Unclassified:    133886     30709277 bp   12.19 %

Total interspersed repeats: 48391211 bp   19.21 %


Small RNA:         1277       772817 bp    0.31 %

Satellites:          25         3445 bp    0.00 %
Simple repeats:     223        23039 bp    0.01 %
Low complexity:       0            0 bp    0.00 %
==================================================
```

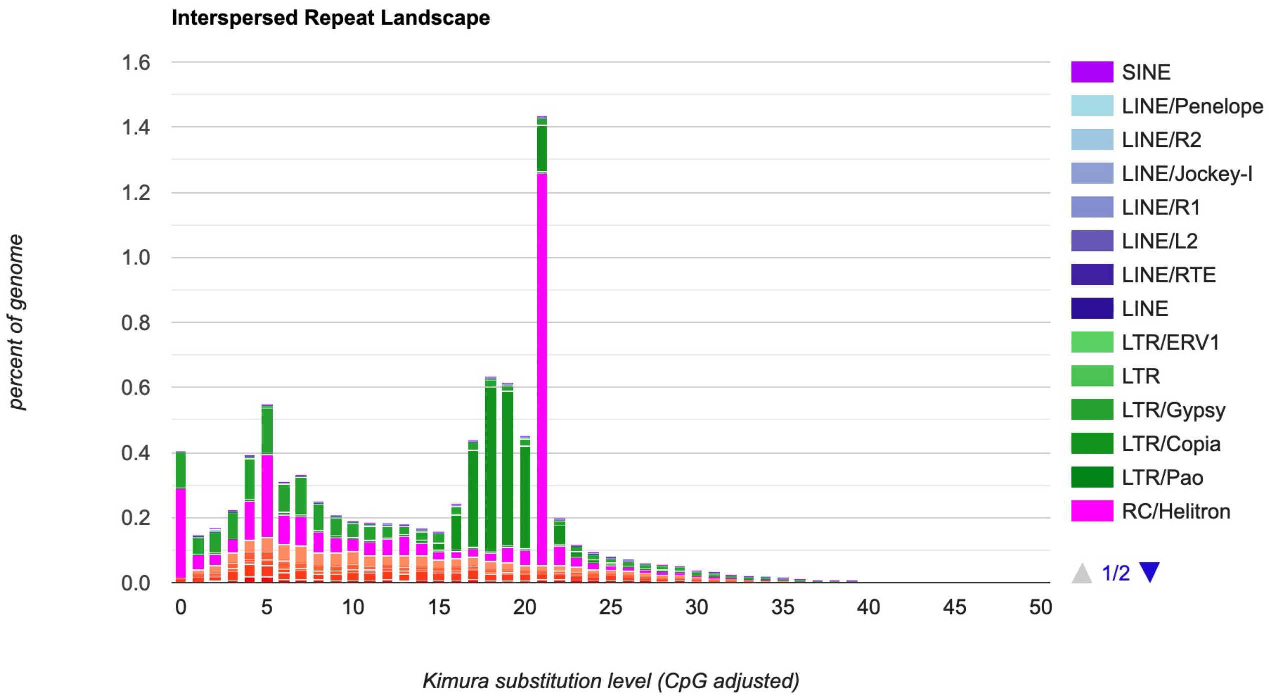


***Figure 4:*** *Kimura distance-based analysis of transposable elements for the genome of P. californicus*

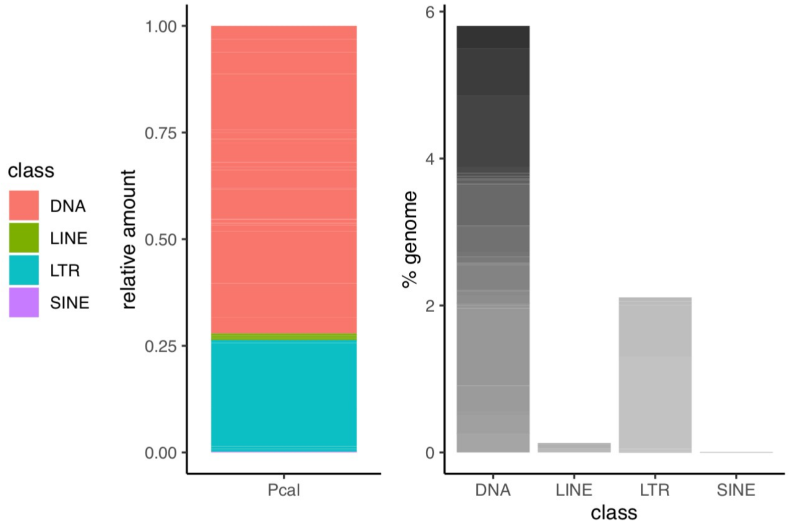


***Figure 5:*** *Summary of transposable element content of the P. californicus genome*

#### Gene annotation

Protein-coding and tRNA genes were annotated with funannotate v1.7.1 in the soft-masked genome. We used short-read Illumina paired-end and long-read minION RNAseq data for training. Homology-based gene predictions were included by running GeMoMa v1.7 on a set of five published ant genomes.

| species | accession |
|---|---|
| Pogonomyrmex barbatus | GCF_000187915.1 |
| Oocecaea biroi | GCF_003672135.1 |
| Camponotus floridanus | GCF_003227725 |
| Solenopsis invicta | GCF_000188075.2 |
| Acromyrmex echinatior | GCF_000204515.1 |

We used paired-end short-read RNAseq data from RNA extracted from heads of 18 individuals of both populations (haplo and pleo) (Ernst et al., unpublished). In total, these accumulated to **9,780,335** read pairs of **1.46 Gbp**.

Funannotate intitially produced 26,408 gene models. After functional annotation with `interproscan` , EggNog, and `funannotate annotate` , we removed any protein-coding gene that did not have any functional annotation or known ortholog and did not have any RNAseq coverage. Of the predicted genes, 9435 had at least some RNAseq support and 15110 showed at least some homology to known proteins (with Evalue < 1E-5 against UniRef90 Release: 2020_04, 12-Aug-2020). 1279 genes had RNAseq support but were not homologous to any protein in UniRef90.

This resulted in an annotation comprising **16,501** genes.

| | **P. californicus genome** |
|---|---|
| **Number of genes** | **16501** |
| **Number of mRNAs** | **17033** |
| Number of exons | 85140 |
| Number of introns | 68107 |
| Number of CDS | 15899 |
| Overlapping genes | 698 |
| Contained genes | 48 |
| mean gene length | 4131 |
| mean mRNA length | 4184 |
| mean exon length | 312 |
| mean intron length | 658 |
| % of genome covered by genes | 27.0 |
| % of genome covered by CDS | 8.8 |
| mean mRNAs per gene | 1 |
| mean exons per mRNA | 5 |
| mean introns per mRNA | 4 |

A `BUSCOv3` (vs. hymenoptera_odb9) analysis of the annotation showed that over 96% of BUSCOs were identified: **C:96.1%[S:90.8%,D:5.3%],F:1.9%,M:2.0%,n:4415**

Functional annotation with InterProScan and EggNog resulted in functional annotations for the following number of genes:

| **Type** | **#Genes** |
|---|---|
| InterPro | 10863 |
| GeneOntology | 7500 |
| PFAM | 9532 |
| EggNog | 6218 |
